# Impact of energy limitations on function and resilience in long-wavelength Photosystem II

**DOI:** 10.1101/2022.04.05.486971

**Authors:** Stefania Viola, William Roseby, Stefano Santabarabara, Dennis Nürnberg, Ricardo Assunção, Holger Dau, Julien Sellés, Alain Boussac, Andrea Fantuzzi, A William Rutherford

## Abstract

Photosystem II (PSII) uses the energy from red light to split water and reduce quinone, an energy-demanding process based on chlorophyll a (Chl-a) photochemistry. Two kinds of cyanobacterial PSII can use Chl-d and Chl-f to perform the same reactions using lower energy, far-red light. PSII from *Acaryochloris marina* has Chl-d replacing all but one of its 35 Chl-a, while PSII from *Chroococcidiopsis thermalis*, a facultative far-red species, has just 4 Chl-f and 1 Chl-d and 30 Chl-a. From bioenergetic considerations, the far-red PSII were predicted to lose photochemical efficiency and/or resilience to photodamage. Here, we compare enzyme turnover efficiency, forward electron transfer, back-reactions and photodamage in Chl-f-PSII, Chl-d-PSII and Chl-a-PSII. We show that: i) all types of PSII have a comparable efficiency in enzyme turnover; ii) the modified energy gaps on the acceptor side of Chl-d-PSII favor recombination via P_D1_^+^Phe^-^ repopulation, leading to increased singlet oxygen production and greater sensitivity to high-light damage compared to Chl-a-PSII and Chl-f-PSII; ii) the acceptor-side energy gaps in Chl-f-PSII are tuned to avoid harmful back reactions, favoring resilience to photodamage over efficiency of light usage. The results are explained by the differences in the redox tuning of the electron transfer cofactors Phe and Q_A_ and in the number and layout of the chlorophylls that share the excitation energy with the primary electron donor. PSII has adapted to lower energy in two distinct ways, each appropriate for its specific environment but with different functional penalties.

## 1 Introduction

Photosystem II (PSII) is the water/plastoquinone photo-oxidoreductase, the key energy converting enzyme in oxygenic photosynthesis. Standard PSII contains 35 chlorophylls a (Chl-a) and 2 pheophytins (Phe). Four of the Chl molecules (P_D1_, P_D2_, Chl_D1_ and Chl_D2_) and both Phe molecules are located in the reaction center (1). The remaining 31 Chl-a in the PSII core constitute a peripheral light-collecting antenna. When antenna chlorophylls are excited by absorbing a photon, they transfer the excitation energy to the primary electron donor, Chl_D1_, the red-most chlorophyll in the reaction center, although it’s been reported that charge separation from P_D1_ can occur in a fraction of centers (1–4). The initial charge separation, forming the first radical pair Chl_D1_^*^ Phe^-^ (assuming Chl_D1_ as primary donor), is quickly stabilized by the formation of the second radical pair, P_D1_^+^ Pheo^-^, and then by further electron transfer steps (Fig. 1A) that lead to the reduction of plastoquinone and the oxidation of water.

**Fig. 1.**
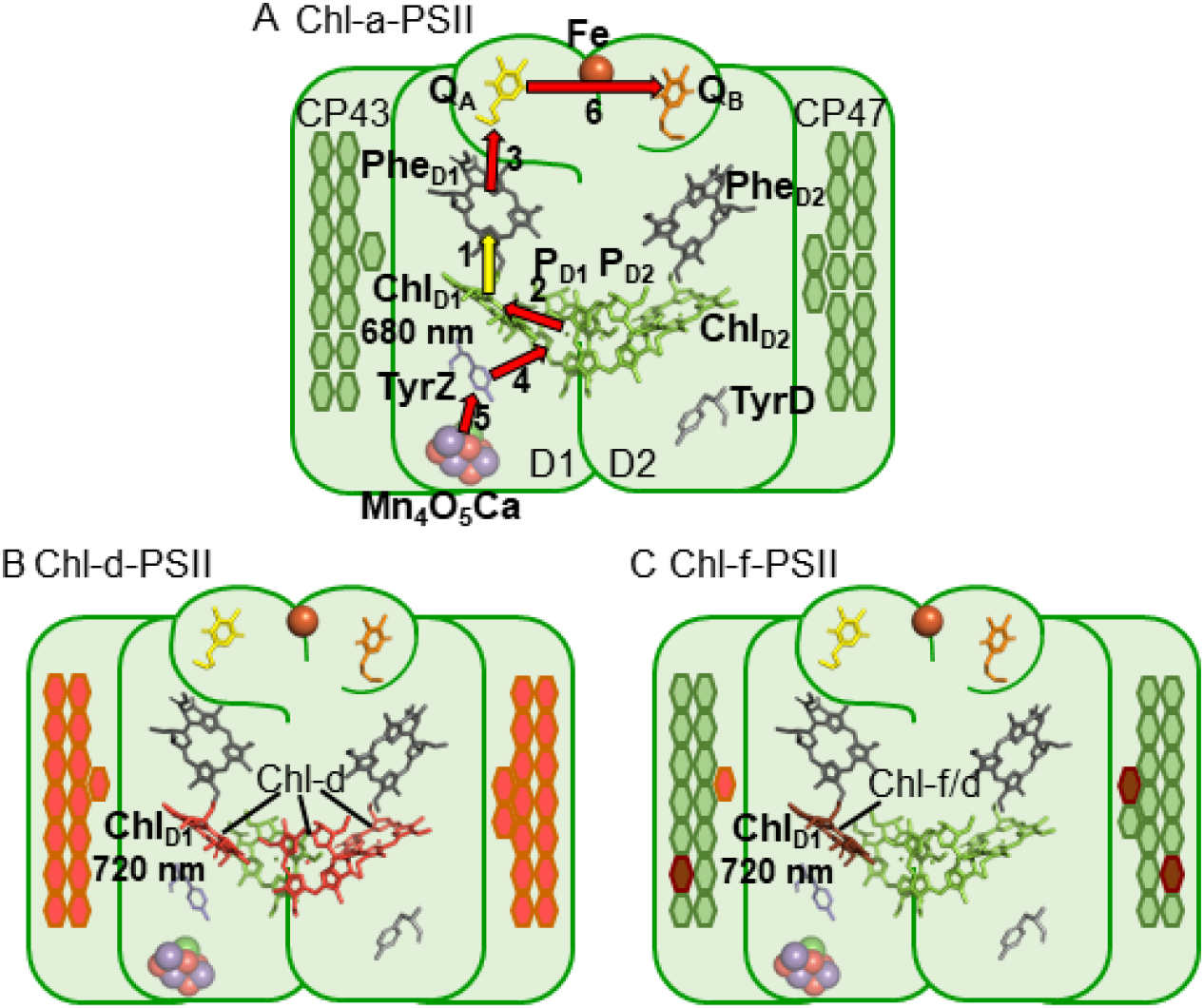
The three types of PSII. (A) Chl-a-PSII (PDB ID: 3ARC, (14)) with the key cofactors of the reaction center, located in the subunits D1 and D2, labelled. Besides the P_D1_, P_D2_, Chl_D1_ and Chl_D2_ chlorophylls and the two pheophytins, Phe_D1_ and Phe_D2_, these cofactors include the quinones, Q_A_ and Q_B_, and the non-heme iron (Fe) on the acceptor side and the two redox-active tyrosines TyrZ and TyrD and the manganese cluster (Mn_4_O_5_Ca) on the donor side. The arrows represent the electron transfer steps and the numbers the order of the steps. The yellow arrow is the primary charge separation, with other steps shown as red arrows. The primary donor is shown as Chl_D1_. (B) and (C) Chl-d-PSII and Chl-f-PSII, with the far-red chlorophylls in the reaction centers highlighted and the wavelength of the primary donor, assumed to be Chl_D1_, indicated. The hexagons on the sides of each reaction center represent the chlorophylls of the respective antennas, located in the subunits CP43 and CP47. Chl-a is represented in green, Chl-d in orange and Chl-f in brown. In (C) the single Chl-d is located in the antenna, but the possibility that it is located in the Chl_D1_ position and plays the primary donor role also exists (5, 13) and the locations of the 4 antenna in the peripheral antenna are uncertain but reflect suggestions in the literature (6).

PSII activity is energy demanding. In Chl-a-PSII, the primary donor absorbs red photons at 680 nm, and this defines the energy available for photochemistry (1.82 eV) with a high quantum yield for the forward reactions. The energy stored in the products of the reaction and in the electrochemical gradient is ∼1 eV, while the remaining ∼0.82 eV is released as heat helping to ensure a high quantum yield for the forward reaction and minimize damaging and wasteful side and back reactions. The 1.82 eV was suggested to be the minimum amount of energy required for an optimum balance of efficiency versus resilience to photodamage, and responsible for explaining the “red limit” (∼680 nm) for oxygenic photosynthesis (5, 6).

The first reported case in which the red limit is exceeded was the Chl-d-containing cyanobacterium *Acaryochloris marina* (*A. marina*) (7). Chl-d-PSII contains 34 Chl-d and 1 Chl-a (proposed to be in the P_D1_ position (8)) and uses less energy, with the proposed Chl-d primary donor in the Chl_D1_ position absorbing far-red photons at ∼720 nm (9), corresponding to an energy of ∼1.72 eV (Fig. 1B).

Recently, it was discovered that certain cyanobacteria use an even more red-shifted pigment, Chl-f, in combination with Chl-a (10, 11). When grown in far-red light, these cyanobacteria replace their standard Chl-a-PSII with Chl-f-PSII, that has far-red specific variants of the core protein subunits (D1, D2, CP43, CP47 and PsbH) and contains ∼90% of Chl-a and ∼10% of Chl-f (5, 11). The Chl-f-PSII from *Chroococcidiopsis thermalis* PCC7203 (*C. thermalis*), which contains 30 Chl-a, 4 Chl-f and 1 Chl-d, was shown to have a long wavelength primary donor (either Chl-f or d, in the Chl_D1_ position) absorbing far-red photons at ∼720 nm (Fig. 1C), the same wavelength as in *A. marina* (5, 12). A recent cryo-EM structure has also argued for Chl_D1_ being the single Chl-d in the far-red PSII of *Synechococcus elongatus* PCC7335 (13). The facultative, long-wavelength species that use Chl-f are thus the second case of oxygenic photosynthesis functioning beyond the red-limit (5), but the layout of their long wavelength pigments is quite different from that of the Chl-d-PSII.

Assuming that Chl-a-PSII already functions at an energy red limit (6), the diminished energy in Chl-d-PSII and Chl-f-PSII seems likely to increase the energetic constraints. Thus, if the far-red PSII variants store the same amount of energy in their products and electrochemical gradient, as seems likely, then it was suggested that they should have decreased photochemical efficiency and/or a loss of resilience to photodamage (5, 15, 16). These predicted energetic constraints are worth investigating to generate knowledge that could be beneficial for designing strategies aimed at engineering of far-red photosynthesis into other organisms of agricultural or technological interest (17).

Here we report a comparison of the enzyme turnover efficiency, forward reactions, and back-reactions in the three known types of PSII: the “standard” Chl-a-PSII, and the two far-red types, the Chl-f-PSII from *C. thermalis* and the Chl-d-PSII from *A. marina*. To compare the enzymatic properties of the three types of PSII and minimize the effects of physiological differences between strains, isolated membranes rather than intact cells were used. The use of isolated membranes allows the minimization of potential effects due to: i) the transmembrane electric field, which affects forward electron transfer (18) and charge recombination (19), ii) the uncontrolled redox state of the plastoquinone pool in whole cells,, which can affect the Q_B_/Q_B_^-^ratio present in dark-adapted PSII, iii) differences in the size and composition of the phycobilisomes and in their association with PSII, and iv) the presence of photoprotective mechanisms such as excitation energy quenching and scavengers of reactive oxygen species.

## 2 Results

### 2.1 Fluorescence decay kinetics in the three types of PSII

The electron transfer properties of the three types of PSII were investigated by comparing the decay kinetics of the flash-induced fluorescence in membranes from *A. marina*, white-light (WL) grown *C. thermalis* and far-red-light (FR) grown *C. thermalis*. When forward electron transfer occurs (Fig. 2A), the fluorescence decay comprises three phases (20, 21): the fast phase (∼0.5 ms) is attributed to electron transfer from Q_A_^-^ to Q_B_ or Q_B_^-^and the middle phase (∼3 ms) is generally attributed to Q_A_^-^oxidation limited by plastoquinone (PQ) entry to an initially empty Q_B_ site and/or by Q_B_H_2_ exiting the site prior to PQ entry (22). These two phases had comparable time-constants in all samples (T_1_ = 0.5-0.6 and T_2_ = 3.5-5 ms, Table S1). The fast electron transfer from Q_A_^-^to the non-heme iron possibly oxidized in a fraction of centers is too fast (t½ ∼50 µs) to be detected here.

**Fig. 2.**
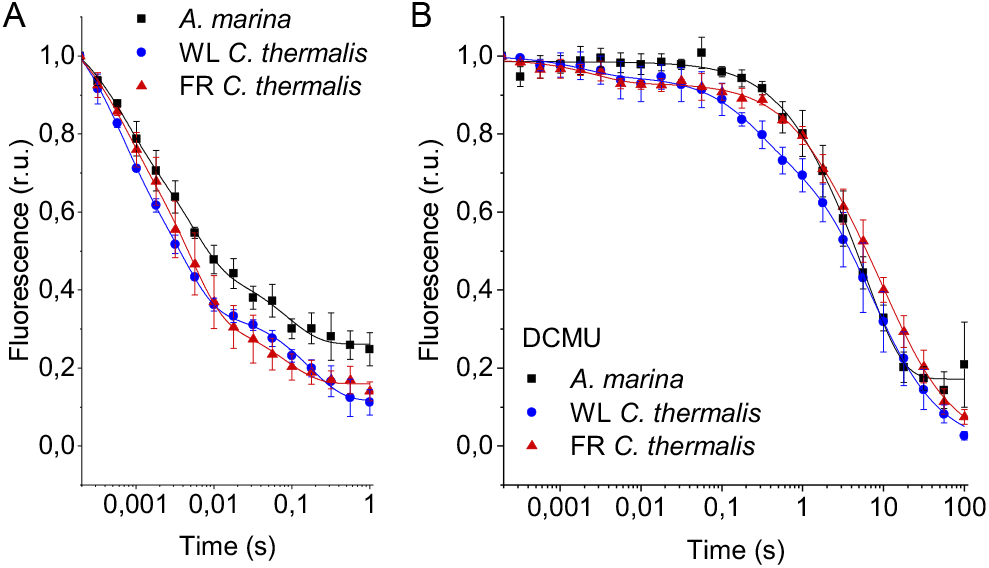
Fluorescence decay kinetics after a saturating flash in membranes of *A. marina*, WL *C. thermalis* and FR *C. thermalis* with no additions (A) and in presence of DCMU (B). The datapoints represent the averages of three biological replicates, ± s.d., the lines represent the fits of the experimental data. All traces are normalized on the initial variable fluorescence (F_m_-F_0_, with F_m_ measured 190 μs after the saturating flash). The full 100 s traces of the data in (A) are shown in Fig. S1.

The slower decay phase is attributed to the charge recombination between Q_A_^-^and the the Mn cluster in the S_2_ state in centers where forward electron transfer to Q_B_/Q_B_^-^did not occur. This phase was significantly slower in FR *C. thermalis* (T_3_ = 14.3±4.6 s) than in WL *C. thermalis* (T_3_ = 5.6±2.4 s) but had a similar amplitude in the two samples (Fig. S1 and Table S2). In *A. marina* this phase had a bigger amplitude than in the two *C. thermalis* samples (Tables S1 and S2), because it was superimposed to a non-decaying component of the fluorescence, that did not return to the original F_0_ level even at 100 s after the flash (Fig. S1). This non-decaying component, absent in the two *C. thermalis* samples, is attributed to centers without a functional Mn-cluster, in which P_D1_^+^ is reduced by an electron donor that does not recombine in the minutes timescale (such as Mn^2+^, TyrD, or the ChlZ/Car side-path), with the consequence of stabilizing Q_A_^-^(23, 24). The fluorescence decay arising from the S_2_Q_A_^-^recombination was slower in *A. marina* (T_3_ = 10.8±2.6 s) than in WL *C. thermalis*, but its overlap with the non-decaying component made the fit of its time-constant potentially less reliable.

Indeed, when the fluorescence decay due to the S_2_Q_A_^-^ recombination was measured in presence of the Q_B_-site inhibitor DCMU (Fig. 2B), the decay kinetics were bi-phasic in all samples, and no difference in the two S_2_Q_A_^-^recombination phases (middle and slow phase in Table S1) was found between *A. marina* and WL *C. thermalis*. In contrast, the decay was significantly slower in FR *C. thermalis*, with the time-constant of the major S_2_Q_A_^-^recombination phase (slow phase in Table S1, ∼80% amplitude, T_3_ = 10.4±0.8 s) similar to that measured in the absence of DCMU (Table S2). The fluorescence decay in WL and FR *C. thermalis* both had an additional fast phase of small amplitude (5-6%), attributed to forward electron transfer in centers in which DCMU was not bound (25). Again, the *A. marina* traces included a non-decaying phase of fluorescence, attributed to centers lacking an intact Mn-cluster.

The fluorescence decay kinetics in membranes of *Synechocystis* sp. PCC6803 (*Synechocystis*), perhaps the best studied Chl-a containing cyanobacterium, were also measured as an additional control. The kinetics in *Synechocystis* membranes were comparable to those reported for WL *C. thermalis* (Fig. S2). The *Synechocystis* and *A. marina* fluorescence decay kinetics measured in membranes here are overall slower than those previously measured in cells (26). This difference is ascribed to pH and membrane potential effects, as discussed in the Supplementary Information, and illustrates the difficulty to use whole cells for such measurements

To conclude, the forward electron transfer rates from Q_A_^-^to Q_B_/Q_B_^-^are not significantly different in the three types of PSII. In contrast, the S_2_Q_A_^-^recombination is slower in Chl-f-PSII of FR *C. thermalis* compared to Chl-a-PSII of WL *C. thermalis* and Chl-d-PSII of *A. marina*.

### 2.2 S-state turnover efficiency in the far-red PSII

The efficiency of PSII activity can be estimated by the flash-dependent progression through the S-states. This can be measured by thermoluminescence (TL), which arises from radiative recombination of the S_2_Q_B_^-^and S_3_Q_B_^-^states (27). The TL measured in *A. marina*, WL *C. thermalis* and FR *C. thermalis* membranes showed similar flash-dependencies in all three types of PSII (Fig. S3), confirming and extending the earlier report (5). Because the TL data presented some variability between biological replicates (see Fig. S3 and associated text), additional analyses were performed by polarography and absorption spectroscopy.

Fig. 3 shows the flash-dependent oxygen evolution measured in *A. marina*, FR *C. thermalis* and *Synechocystis* membranes. The latter were used as a Chl-a-PSII control because the content of PSII in membranes of WL *C. thermalis* was too low to allow accurate O_2_ polarography measurements. As shown by fluorescence (Fig. S2), no significant difference in forward electron transfer between the two types of Chl-a-PSII was observed, and the use of *Synechocystis* membranes was therefore considered as a valid control.

**Fig. 3.**
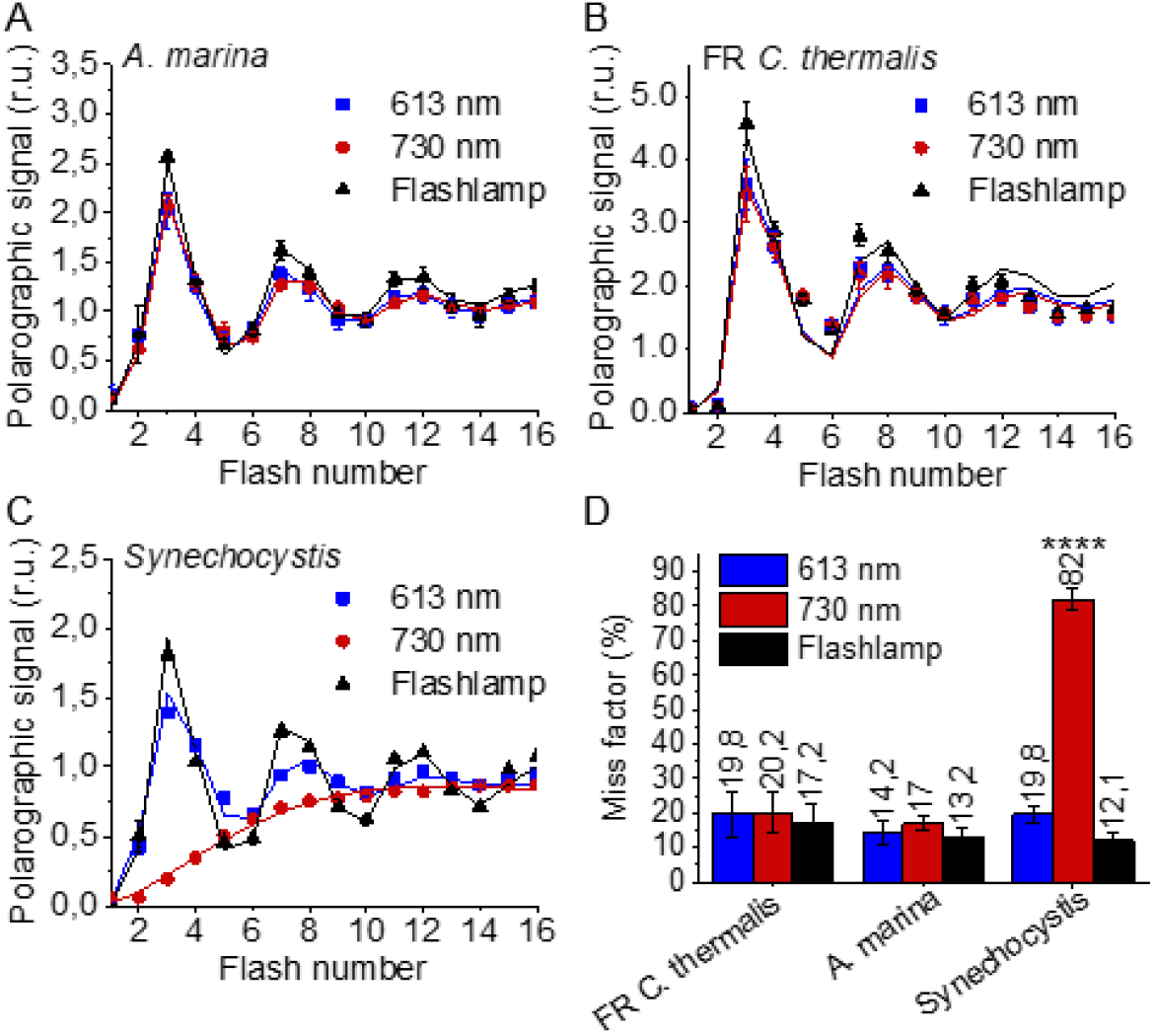
Flash-induced release of O_2_ measured by polarography. (A-C) Patterns of oxygen release in *A. marina*, FR *C. thermalis* and *Synechocystis* membranes. Flashes were given at 900 ms intervals and the O_2_ produced after each flash was measured. Flashes were provided by a white xenon flash lamp, a red LED centered at 613 nm, and a far-red LED centered at 730 nm. The data represent the averages of 3 biological replicates ±s.d. The lines represent the fits of the experimental data. (D) Miss factors (in %) calculated from the data shown in (A-C). The miss factor in *Synechocystis* membranes flashed at 730 nm is significantly higher than in *A. marina* and FR *C. thermalis* membranes according to Student’s t-test, as indicated with asterisks (****p≤ 0.0001).

The measurements were performed using white, red, and far-red flashes. As expected, in dark-adapted samples, with S_1_ as the majority state, the maximal O_2_ evolution occurred on the 3^rd^ flash with subsequent maxima at 4 flash intervals. These maxima reflect the occurrence of the S_3_Y_Z_ ^●^/S_4_ to S_0_ transition in most centers as two water molecules are oxidized, resulting in the release of O_2_. This oscillation pattern was the same in all samples and under all excitation conditions, except in *Synechocystis* membranes illuminated with far-red light, where the slow rise in O_2_ evolution is due to the weak excitation of Chl-a-PSII by the short wavelength tail of the 730 nm flash.

The miss factor (Fig. 3D) was ≤20% in all the samples except in the *Synechocystis* sample illuminated with far-red flashes (>80%). For *A. marina*, the misses (13-17%) were very similar to those reported earlier (28). The misses in FR *C. thermalis* and in *Synechocystis* when illuminated with the 613 nm LED were slightly higher (17-20%). Nevertheless, these differences, attributed to the combination of the absence of exogenous electron acceptors, and the relatively long and possibly not fully saturating flashes, were not significant.

In order to confirm and expand the results obtained with polarography, we measured the S-state turnover as the flash-induced absorption changes at 291 nm (Fig. 4), that reflect the redox state of the Mn ions in the oxygen evolving complex (29). These measurements were done in the presence of the electron acceptor PPBQ and using single-turnover monochromatic saturating laser flashes. In the case of *A. marina*, the measurements could be done using membranes, but the membranes of WL and FR *C. thermalis* could not be used because of their high light-scattering properties in the UV part of the spectrum. In the case of the FR *C. thermalis* partially purified O_2_ evolving Chl-f-PSII were made and used for the measurements, while difficulties were encountered in isolating O_2_ evolving PSII from WL *C. thermalis*. Therefore, PSII cores from *T. elongatus* with the D1 isoform PsbA3, which has the highest sequence identity with the D1 of Chl-f-PSII in FR *C. thermalis* (see discussion), were used as a Chl-a-PSII control (30).

**Fig. 4.**
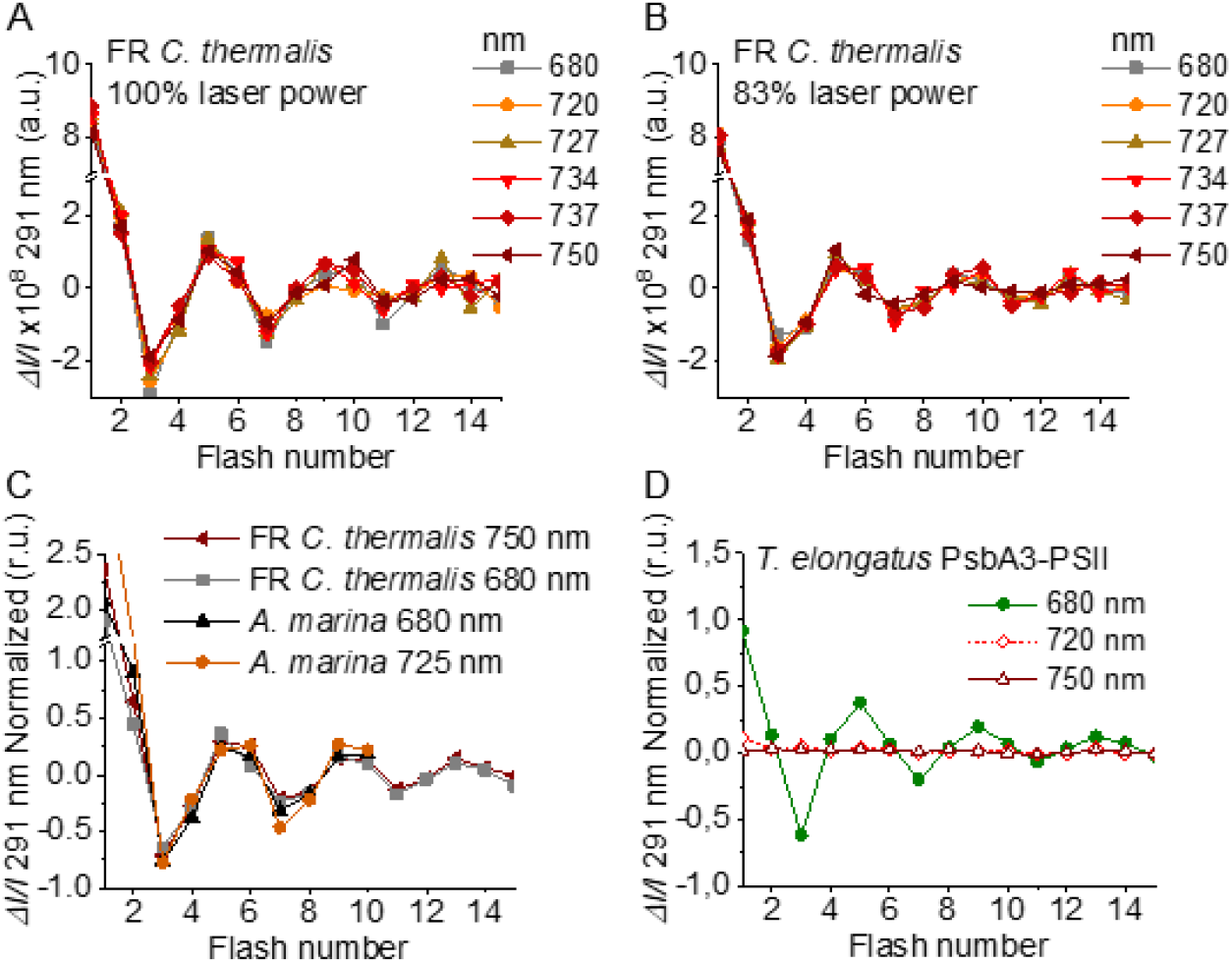
Flash-induced S-state turnover in FR *C. thermalis* PSII cores, *A. marina* membranes, and *T. elongatus* PsbA3-PSII cores. Absorption changes were measured at 291 nm at 100 ms after each of a series of single-turnover saturating flashes fired with a 300 ms time interval. (A) and (B) Measurements in FR *C. thermalis* PSII cores using flashes at the indicated wavelengths with 100% and 83% laser power (the power of the laser at the different wavelengths is reported in the Supplementary Materials and Methods). (C) Comparison between the absorption changes obtained in FR *C. thermalis* PSII cores and *A. marina* membranes using flashes at the indicated wavelengths (100% laser power). The traces in (C) were normalized on the maximal oscillation amplitude (3^rd^ minus 5^th^ flash). The breaks in the vertical axes in panels (A-C) allow the oscillation pattern to be re-scaled for clarity, because the absorption change on the first flash contains a large non-oscillating component (29) that was not included in the fits. (D) Measurements in isolated *T. elongatus* PsbA3-PSII cores using flashes at the indicated wavelengths.

The Chl-f-PSII was illuminated with flashes at wavelengths preferentially absorbed by Chl-a (680 nm) and of the long-wavelength chlorophylls (720 to 750 nm) (Fig. 4A). As expected, maximum Δ*I/I* occurred on S_2_ (flash 1,5,9 etc.) and minimum Δ*I/I* on S_0_ (flash 3,7,11 etc.) (29). No differences could be observed in either the amplitude or the damping of the oscillations between the excitation wavelengths. When using sub-saturating flashes (∼83% power), the damping of the oscillations was the same for all excitation wavelengths (Fig. 4B), verifying that the illumination with 100% laser power was saturating at all the wavelengths. The equal amplitude of the oscillations obtained at all excitation wavelengths also indicates that the FR *C. thermalis* sample used does not contain any detectable Chl-a-PSII contamination. No differences in the oscillation patterns measured in FR *C. thermalis* Chl-f-PSII cores and in *A. marina* membranes, flashed at either 680 or 725 nm, were observed (Fig. 4C). The PSII of *T. elongatus* showed a normal S-states progression when using 680 nm excitation, but no oscillation pattern when far-red flashes were used (Fig. 4D). For all samples the calculated miss factor was ∼10% (Fig. S4).

In conclusion, the data reported here show that the overall efficiency of electron transfer from water to the PQ pool is comparable in all three types of PSII (independently of the Chl-a-PSII control used), as shown by the near-identical flash patterns of thermoluminescence (Fig. S3) and O_2_ release (Fig. 3), both measured without external electron acceptors. When the S-state turnover was measured by following the absorption of the Mn cluster in the UV (Fig. 4), the use of artificial electron acceptors and single-turnover saturating flashes allowed us to obtain better resolved flash patterns that were essentially indistinguishable in all three types of PSII and between excitation with visible or far-red light in the case of the Chl-d-PSII and Chl-f-PSII.

### 2.3 Back-reactions measured by (thermo)luminescence

Charge recombination reactions were investigated by monitoring the thermoluminescence and luminescence emissions. The TL curves in Fig. 5A and B show that both Chl-f-PSII and Chl-d-PSII are more luminescent than Chl-a-PSII, with Chl-f-PSII being the most luminescent. These differences, that are much larger than the variability between biological replicates (Fig. S5 and Table S3), fit qualitatively with earlier reports (5, 31). The high luminescence indicates that in the Chl-d-PSII and Chl-f-PSII there is an increase in radiative recombination, although the causes of this increase are likely to be different between the two photosystems, as detailed in the Discussion.

**Fig. 5.**
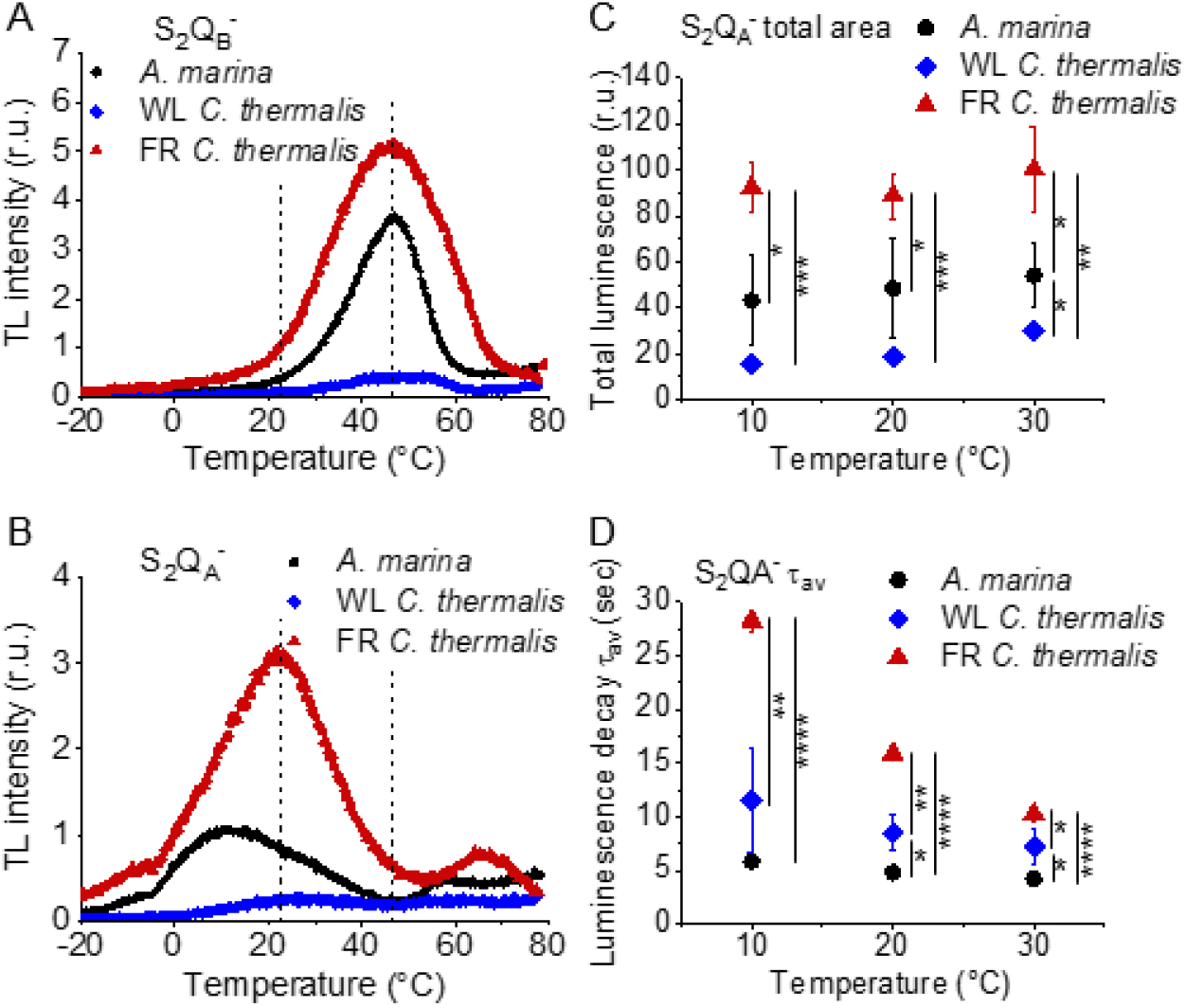
Thermoluminescence and luminescence measured in *A marina*, WL *C. thermalis* and FR *C. thermalis* membranes. (A) and (B) TL measured in the absence of inhibitors (S_2_Q_B_^-^) or in the presence of DCMU (S_2_Q_A_^-^), respectively. The signal intensities are normalized on the maximal oxygen evolution rates of each sample. The dashed vertical lines indicate the two peak positions of the *C. thermalis* samples. (C) Plots of the total S_2_Q_A_^--^ luminescence emission (integrated area below the curves), normalized on the maximal oxygen evolution rate of each sample, at 10, 20 and 30°C. (D) Plots of the average S_2_Q_A_^--^ luminescence decay lifetimes (τ_av_), calculated from the decay phases attributed to S_2_Q_A_^--^ recombination, as a function of temperature. In (C) and (D) each point represents the average of 3 biological replicates ±s.d. Statistically significant differences according to Student’s t-tests are indicated with asterisks (*p ≤ 0.05, **p ≤ 0.01, ***p ≤ 0.001, ****p≤ 0.0001).

Despite the large difference in TL intensity between the Chl-a-PSII and Chl-f-PSII, the peak temperatures corresponding to the S_2_Q_B_^-^and S^2^Q_A_^-^ recombination were both similar in Chl-a-PSII and Chl-f-PSII. In Chl-d-PSII, the temperature of the S_2_Q_B_^-^ peak was only slightly lower, while the S_2_Q_A_^-^peak was ∼15°C lower (Fig. S5 and Table S3). Earlier TL reports comparing Chl-d-PSII in *A marina* cells with Chl-a-PSII in *Synechocystis* cells also showed that, while the peak position of S_2_Q_B_^-^ recombination was similar in the two samples, the S_2_Q_A_^-^ peak position was lower in *A. marina* (31), in agreement with the present results in membranes. The peak temperatures measured in cells were lower than those reported here, which can be explained by i) the effect of the transmembrane electric field, as discussed for the fluorescence decay (section 2.1), and ii) by differences in the heating rates used (1°C s^-1^ here, 0.33°C s^-1^ in (31)). When performing the same measurements in *Synechocystis* membranes (Fig. S6), the S_2_Q_B_^-^ and S_2_Q_A_^-^ peak positions were comparable to those obtained in the two *C. thermalis* samples, confirming that the lower S_2_Q_A_^-^ peak temperature is a specific feature of Chl-d-PSII.

The S_2_Q_A_^-^ recombination in the presence of DCMU was also measured by luminescence decay kinetics at 10, 20 and 30°C, a range of temperatures that covers those of the S_2_Q_A_^-^ TL peaks of the three samples. Luminescence decay kinetics were recorded from 570 ms for 300 seconds after the flash. In this time-range, the luminescence arises mainly from recombination *via* the back-reaction of S_2_Q_A_^-^ (32). The total S_2_Q_A_^-^ luminescence emission (Fig. 5C) reflected the intensities of the TL peaks, as expected (33), with the order of intensity as follows: Chl-f-PSII > Chl-d-PSII > Chl-a-PSII (although the variability between replicates made the difference between Chl-a-PSII and Chl-d-PSII less significant than that measured by TL). The total emissions did not vary significantly between 10 and 30°C, although the decay kinetics were temperature-sensitive (Fig. S7). The decay components identified by fitting the curves and their significance are discussed further in the SI. The luminescence decay attributed to S_2_Q_A_^-^ recombination was bi-phasic (Table S4), with the kinetics of both phases being faster in Chl-d-PSII (∼3 and ∼11 s) than in Chl-a-PSII (∼4 and ∼25 s), but slower in Chl-f-PSII (∼9 and ∼39 s). The average S_2_Q_A_^-^ luminescence decay lifetimes accelerated with increasing temperature in Chl-a-PSII and Chl-f-PSII but were always the fastest in Chl-d-PSII and the slowest in Chl-f-PSII (Fig. 5D). The luminescence decay kinetics of the Chl-a-PSII in *Synechocystis* membranes were similar to those measured in WL *C. thermalis* (Fig. S8), suggesting, as seen with the TL data, that the differences in kinetics observed in the two types of far-red PSII are not due to differences between species.

In conclusion, both Chl-f-PSII and Chl-d-PSII show strongly enhanced luminescence, as previously reported (5, 34). However, the Chl-d-PSII differs from the Chl-a-PSII and Chl-f-PSII by having a lower S_2_Q_A_^-^ TL peak temperature and a faster S_2_Q_A_^-^ luminescence decay. This indicates that Chl-d-PSII has a smaller energy gap between Q_A_^-^ and Phe compared to Chl-a-PSII and Chl-f-PSII. The lower TL temperature and faster luminescence decay for S_2_Q_A_^-^ recombination in Chl-d-PSII, but without a marked increase in its Q_A_^-^ decay rate as monitored by fluorescence (Fig. 2), could reflect differences in the competition between radiative and non-radiative recombination pathways in Chl-d-PSII compared to those in Chl-a-PSII and Chl-f-PSII. In contrast, in Chl-f-PSII the energy gap between Q_A_^-^ and Phe does not appear to be greatly affected or could even be larger, as suggested by the slower S_2_Q_A_^-^ recombination measured by fluorescence (Fig. 2) and slower luminescence (Fig. 3) decay. The Q_B_ potentials appear to be largely unchanged, as manifest by the similar S_2_Q_B_^-^ stability in all three types of PSII, with the slightly lower S_2_Q_B_^-^ TL peak temperature in *A. marina*, probably reflecting the decrease in the energy gap between Q_A_^-^ and Phe.

## 2.4 Singlet oxygen production and sensitivity to high light in the far-red PSII

The smaller energy gap between Q_A_^-^ and Phe reported here in *A marina* is expected to result in enhanced singlet O_2_ production and hence greater sensitivity to photodamage (5, 15, 35, 36). This was investigated by measuring the rates of ^1^O_2_ generation induced by saturating illumination in isolated membranes using histidine as a chemical trap (Fig. 6A, representative traces in Fig. S10A-C). ^1^O_2_ reacts with histidine to form the final oxygenated product, HisO_2_, resulting in the consumption of O_2,_ as measured using the O_2_ electrode. Without the histidine trap, most ^1^O_2_ is thought to be quenched by carotenoids (37). When histidine was present in addition to DCMU, the Chl-d-PSII in *A. marina* membranes showed significant light-induced ^1^O_2_ formation. Under the same conditions, little ^1^O_2_ formation occurred in Chl-a-PSII or Chl-f-PSII in *C. thermalis* membranes. Similarly low levels of ^1^O_2_ were generated by Chl-a-PSII in *Synechocystis* membranes (Fig. S10D). Sodium azide, a ^1^O_2_ quencher, suppressed the His-dependent oxygen consumption measured in the presence of DCMU, confirming that it was due to the production of ^1^O_2_ (Fig. S10E).

**Fig. 6.**
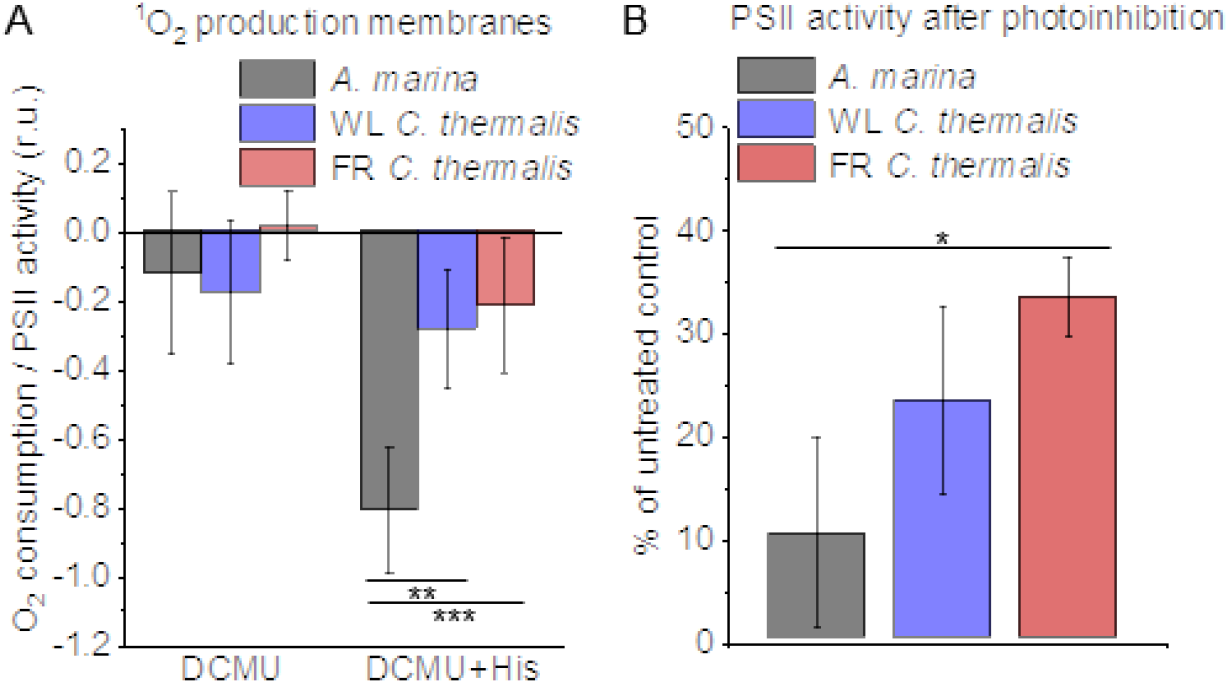
^1^O_2_ production and PSII sensitivity to high light in *A. marina*, WL *C. thermalis* and FR *C. thermalis* membranes. All samples were used at a chlorophyll concentration of 5 µg ml^-1^. (A) ^1^O_2_ production in presence of DCMU measured as the rate of histidine-dependent consumption of O_2_ induced by saturating illumination (xenon lamp, 7100 µmol photons m^-2^ s^-1^, saturation curves in Fig. S9B and C). The data are averages (±s.d.) of 6 biological replicates for *A. marina* and FR *C. thermalis* and 3 replicates for WL *C. thermalis*. For each replicate, the rates of oxygen consumption were normalized to the maximal oxygen evolution rates measured in presence of DCBQ and ferricyanide. (B) Maximal PSII activities, measured as in (A), after 30 min illumination with saturating red light (660 nm LED, 2600 µmol photons m^-2^ s^-1^) relative to the maximal activities measured in control samples kept in darkness. The light used for the 30 minutes treatment was as saturating as the xenon lamp used in (A) (Fig. S9D and E). The data are averages of 3 biological replicates ±s.d. Statistically significant differences according to Student’s t-tests are indicated with asterisks (*p ≤ 0.05, **p ≤ 0.01, ***p ≤ 0.001).

The strikingly high amount of ^1^O_2_ generated by Chl-d-PSII prompted us to perform additional controls. i) To test if the high ^1^O_2_ production was related to the intactness of the PSII donor side, Mn was removed from *A. marina* membranes by Tris-washing. This had little effect on the ^1^O_2_ formation with respect to the Mn-containing membranes (Fig. S11), suggesting that the high ^1^O_2_ production in untreated *A. marina* membranes does not arise specifically from the fraction of centers lacking an intact Mn-cluster that are likely possibly responsible for the non-decaying fluorescence observed in Fig. 2B and S1. ii) The possibility that photosystem I (PSI) contributed to the light-induced O_2_ consumption by reducing oxygen to O_2_^•-^in membranes was tested (Fig. S12). In the presence of DCMU, PSI-driven O_2_ reduction mediated by methyl viologen only took place when exogenous electron donors to PSI were provided. This indicates that there is no contribution from PSI-induced O_2_ reduction in Fig. 4A, where exogenous PSI donors are absent. iii) The higher ^1^O_2_ production is also seen in *A. marina* cells (Fig. S13A) compared to FR *C. thermalis* cells, and thus is not an artefact associated with the isolation of membranes. WL *C. thermalis* cells also showed low levels of ^1^O_2_ production, similar to those measured in membranes (Fig. S13B).

Figure 4B shows the effect of 30 minutes of saturating illumination (red light) on the activity of the Chl-d-PSII, Chl-a-PSII and Chl-f-PSII. The results show that Chl-d-PSII is significantly more susceptible to light induced loss of activity compared to Chl-f-PSII, and to a lesser extent to Chl-a-PSII, and this can be correlated to the higher levels of ^1^O_2_ production in Chl-d-PSII.

## 3 Discussion

We investigated several functional properties of the two different types of far-red PSII, i) the constitutive Chl-d-PSII of *A. marina*, and ii) the facultative Chl-f-PSII of *C. thermalis*. We compared these properties with each other and with those of standard Chl-a-PSII, from either WL *C. thermalis, Synechocystis* or *T. elongatus*, looking for differences potentially related to the diminished energy available in the two long-wavelength PSII variants.

### 3.1 Forward electron transfer and enzymatic activity

The turnover of the water oxidation cycle is comparably efficient in all three types of PSII, as shown by their near-identical flash patterns in thermoluminescence (Fig. S3), O_2_ release (Fig. 3), and UV spectroscopy (Fig. 4). In PSII, a photochemical “miss factor” can be calculated from the damping of the flash patterns of O_2_ evolution. These misses, which are typically ∼10% in Chl-a-PSII, are mainly ascribed to the µs to ms recombination of S_2_TyrZ•Q_A_^-^ and S_3_TyrZ•Q_A_^-^ states (38). Despite the diminished energy available, the miss factors in both types of far-red PSII were virtually unchanged compared to Chl-a-PSII, which also suggests that they have the same origin. If so, the energy gaps between TyrZ and P_D1_, and thus their redox potentials, would be essentially unchanged. These conclusions agree with those in earlier work on Chl-d-PSII (39) and on Chl-f-PSII (5).

The similar flash-patterns also indicate that, after the primary charge separation, the electron transfer steps leading to water oxidation must have very similar efficiencies in all three types of PSII, i.e. close to 90%, and that there are no major changes affecting the kinetics of forward electron transfer, as pointed out earlier based on less complete data (5). Indeed, electron transfer from Q_A_^-^ to Q_B_/Q_B_^-^, monitored by fluorescence, showed no significant differences in kinetics in the three types of PSII (Fig. 2A).

### 3.2 Back reactions and singlet oxygen production

The most striking difference between the three types of PSII is that the Chl-d-PSII of *A. marina* shows a decreased stability of S_2_Q_A_^-^, indicated by the lower temperature of its TL peak and the correspondingly faster luminescent decay kinetics (Fig. 5), and consequently a significant increase in ^1^O_2_ generation under high light (Fig. 6A). This likely corresponds to the decrease in the energy gap between Phe and Q_A_ predicted to result from the ∼100 meV lower energy available when using light at ∼720 nm to do photochemistry (5, 15). This is also supported by the estimates in the literature of the redox potential (E_m_) values of Phe/Phe^-^ and Q_A_/Q_A_^-^ in Mn-containing Chl-d-PSII: compared to Chl-a-PSII, the estimated increase of ∼125 mV in the E_m_ of Phe/Phe^-^ is accompanied by an estimated increase of only ∼60 mV in the E_m_ of Q_A_/Q_A_^-^, which implies that a normal energy gap between the excited state of the primary donor (Chl_D1_^*^) and the first and second radical pairs (Chl_D1_^*^ Phe^-^ and P_D1_^+^ Phe^-^) is maintained, but the energy gap between P_D1_^+^ Phe- and P_D1_^+^ Q_A_^-^ is significantly decreased (∼325 meV vs ∼ 385 meV) (40). The changes in the D1 and D2 proteins of *A. marina* responsible for the changes in the E_m_ of Phe/Phe^-^ and Q_A_/Q_A_^-^ are currently unknown. Our results indicate that in Chl-d-PSII, the decrease in the energy gap between Phe and Q_A_ favors charge recombination by the back-reaction route (via P_D1_^+^ Phe^-^), forming the reaction center chlorophyll triplet state (41), which acts as an efficient sensitizer for ^1^O_2_ formation (35, 36, 42, 43). Consequently, the Chl-d-PSII is more sensitive to high light (Fig. 6B). The increase in the proportion of recombination going via P_D1_^+^ Phe^-^ can also result in a higher repopulation of the excited state of the primary donor (Chl_D1_^*^), with a consequent increase in radiative decay (high luminescence).

In contrast to the Chl-d-PSII, the Chl-f-PSII shows no increased production of ^1^O_2_ and no increased sensitivity to high light compared to Chl-a-PSII, in the conditions tested here (Fig. 6). The back-reactions appear to be little different from the Chl-a-PSII except for the more stable (more slowly recombining) S_2_Q_A_^-^, as seen by fluorescence (Fig. 2) and luminescence (Fig. 5) decay. These properties may seem unexpected because this type of PSII has the same energy available for photochemistry as the Chl-d-PSII. In the Chl-d-PSII the lower energy of Chl_D1_^*^ is matched by an increase in the E_m_ of Phe/Phe^-^. In the Chl-f-PSII of *C. thermalis* and of the other Chl-f containing species, the E_m_ of Phe/Phe^-^ is also expected to be increased by the presence, in the far-red D1 isoform, of the strong H-bond from Glu130 (Fig. S14), which is characteristic of high-light D1 variants in cyanobacteria (44). In Chl-a-PSII this change has been reported to induce an increase in the E_m_ of Phe/Phe^-^ between ∼15 and ∼30 mV (44, 45): an increase of this size would only partially compensate for the ∼100 meV decrease in the energy of Chl_D1_^*^ in Chl-f-PSII, and this would result in a smaller energy gap between Chl_D1_^*^ and the first and second radical pairs Chl_D1_^*^Phe^-^ and P ^+^Phe^-^. This would favor the repopulation of Chl_D1_^*^ by back-reaction from P_D1_^+^Phe^-^ (even if the repopulation of P_D1_^+^Phe^-^ from the P_D1_^+^Q_A_^-^ state did not increase), resulting in the higher luminescence of Chl-f-PSII, as proposed earlier (5).

The longer lifetime of S_2_Q_A_^-^ recombination in Chl-f-PSII indicates that the E_m_ of Q_A_/Q_A_^-^ has increased to compensate the up-shift in the E_m_ of Phe/Phe^-^ and to maintain an energy gap between Phe and Q_A_ large enough to prevent an increase in reaction center chlorophyll triplet formation. This situation occurs in the PsbA3-D1 high light variant of *T. elongatus*, although the protein changes responsible for the increase in the E_m_ of Q_A_/Q_A_^-^ are not known (44). A slower S_2_Q_A_^-^ recombination could also arise from an increase in the redox potential of P_D1_/P_D1_^+^ (46, 47), but this would likely compromise forward electron transfer in Chl-f-PSII by decreasing the driving force for stabilization of Chl_D1_^*^Phe^-^ into P_D1_^+^Phe^-^, if the redox potential of Chl_D1_/Chl_D1_^*^ was not increased accordingly, or by decreasing the already diminished reducing power of Chl_D1_^*^, if the redox potential of Chl_D1_/Chl_D1_^*^ was increased accordingly, which is not what we observe (Fig. 2A).

### 3.5 Effects of the pigment composition on the energetics of the far-red PSII

In addition to changes in the redox potentials of Phe and Q_A_, the size and pigment composition of the antennas of Chl-d-PSII and Chl-f-PSII could also contribute to the functional differences reported in the present work. These differences are summarized in Fig. 7.

**Fig. 7.**
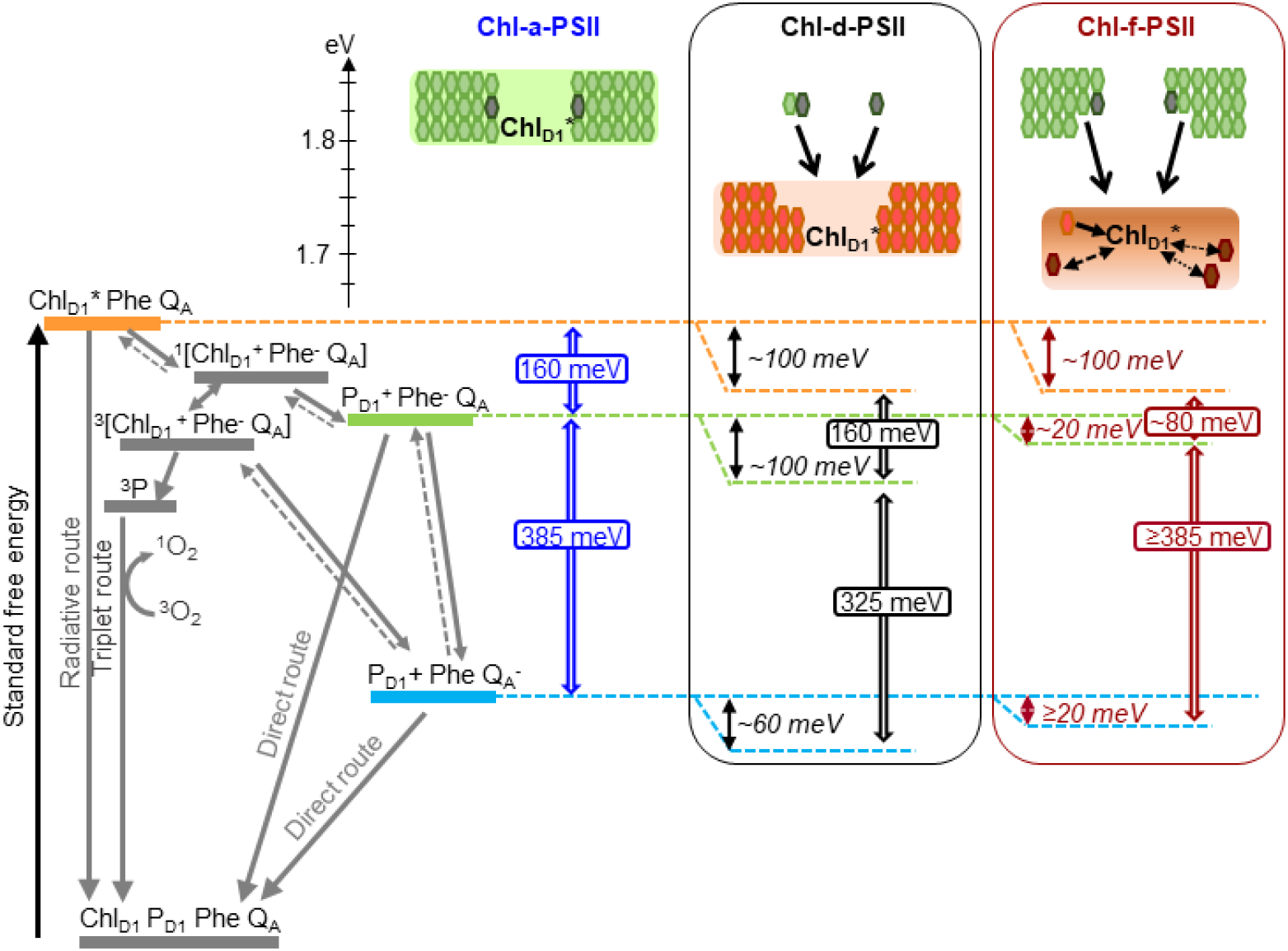
Model of the energy differences in Chl-a-PSII, Chl-d-PSII and Chl-f-PSII. The top part of the figure represents the localization of the excitation energy over the antenna pigments and Chl_D1_* (energies in eV, scale on the left side). The localization of the excitation energy is indicated by the colored boxes (green for Chl-a, orange for Chl-d and brown for Chl-f), without necessarily assuming a full equilibration (see main text). In Chl-a-PSII, the excitation is distributed over Chl_D1_, 34 antenna Chl-a (light green) and 2 Phe-a (dark grey); in Chl-d-PSII, the excitation is distributed over Chl_D1_ and 31 antenna Chl-d (orange) but not over the 1 Chl-a and 2 Phe-a, that transfer excitation downhill to the Chl-d pigments (black arrows); in Chl-f-PSII, the excitation is distributed only over Chl_D1_, one Chl-d and 3 Chl-f (brown), while the remaining 29 Chl-a and 2 Phe-a transfer excitation energy downhill to the far-red pigments. In Chl-f-PSII, 3 of the far-red antenna pigments are at longer wavelength than Chl_D1_, so transfer of excitation energy from them to Chl_D1_ is less efficient (dashed and dotted black arrows). The grading of the colored box for Chl-f represents uncertainty in the degree of excited state sharing between the longest wavelength chlorophylls and Chl_D1_. The bottom part of the figure represents, on the left, the energetics of the radical pairs and the recombination routes in PSII. The horizontal dashed lines represent the standard free energies of Chl_D1_^*^(orange), P_D1_^+^ Phe^-^ (light green) and P_D1+_Q_A_^*^ (light blue). The free energy gaps between Chl_D1_^*^ and P_D1_^+^ Phe^-^ and between P_D1_^+^ Phe^-^ and P_D1_^+^ Q_A_^*^ in Chl-a-PSII (blue) and our current estimates for Chl-d-PSII (black) and Chl-f-PSII (dark red) are represented on the right.

In PSII, two factors will determine the yield of charge separation: i) the relative population of the excited state of the primary donor, Chl_D1_^*^, which depends on the dynamics of excitation energy transfer between pigments, and ii) the rate of population of the second radical pair, P_D1_^+^ Phe^-^, that is more stable (less reversible) than the first radical pair, Chl_D1_^*^ Phe^-^. This rate is determined by the rates of the primary charge separation (forming Chl_D1_^*^ Phe^-^) and of its stabilization by secondary electron transfer (forming P_D1_^+^ Phe^-^), and hence by the energetic of these electron transfer steps.

In the Chl-a-PSII core, the 37 chlorins absorb light between ∼660 and ∼690 nm and are therefore almost isoenergetic to the Chl_D1_ primary donor absorbing at 680 nm. Given the small energy differences, there is little driving force for downhill “funneling” of excitation energy to Chl_D1_, making it a “shallow trap”. Different models have been proposed to explain the shallowness of the photochemical trap in Chl-a-PSII.

In the trap-limited model, the transfer of excitation between pigments is significantly faster than the electron transfer reactions leading to P_D1_^+^Phe^-^ formation, and a near-complete equilibration of the excitation energy is established over all pigments, including Chl_D1_, with a distribution that is determined by their individual site energies (48–50). This leads to a low population of Chl_D1_^*^ (see Table S5), that is diminished as a function of the number of quasi-isoenergetic pigments with which it shares the excitation energy.

In the transfer-to-trap limited model, the small driving force for downhill “funneling” of excitation energy to ChlD1 causes kinetic bottlenecks for excitation energy equilibration between the core antenna complexes CP43 and CP47 and for excitation energy transfer from these antennas to the reaction center (51–54). In this scenario, there is not a full equilibration of the excitation energy over all pigments, but the relatively slow and reversible energy transfer from the core antennas to the reaction center still leads to a relatively low population of Chl_D1_^*^.

Irrespectively of the differences in the details of the kinetic limitation to photochemical trapping between the two models, the common requirement for establishing a high quantum yield of charge separation is a sufficiently large overall energy gap (∼160 meV, (47)) between Chl_D1_* and P ^+^Phe^-^, i.e. comprising the primary charge separation (Chl_D1_^*^ ↔ Chl_D1_^*^Phe^-^) and secondary electron transfer (Chl_D1_^*^Phe^-^ ↔ P_D1_^+^Phe^-^), as shown in Fig. 7. An energy gap of this magnitude is required to avoid rapid recombination to the excited state Chl_D1_^*^, thereby limiting the probability of its dissipation via non-photochemical relaxation to the ground state in the antenna (53, 55).

For Chl-d-PSII the antenna system is comparable to that in Chl-a-PSII: all 34 Chl-d molecules, including the primary donor Chl_D1_ at ∼720 nm, are close in wavelength and thus both systems are expected to have comparable Chl_D1_^*^ population (Table S5), irrespective of the rate-limitation model assumed. Chl-a-PSII and Chl-d-PSII should therefore have the same energetic requirements to ensure a sufficiently high yield of charge separation. Given that the energy of Chl_D1_^*^ is ∼100 meV lower in Chl-d-PSII than in Chl-a-PSII, the energy level of the second and more stable radical pair, P_D1_^+^Phe^-^, needs to be decreased by ∼100 meV in Chl-d-PSII relative to Chl-a-PSII. This corresponds to the published E_m_ of Phe/Phe^-^ (40) and to the kinetic data (Fig. 5 and 6), as detailed in the previous section.

In *A. marina* membranes, additional Chl-d containing antenna proteins, which form supercomplexes with PSII cores, have been reported to increase the Chl-d-PSII antenna size by almost 200% (56). This will likely result in an increased sharing of the excited state, leading to a diminished population of Chl_D1_^*^, and thus a bigger requirement for an energy drop between Chl_D1_^*^ and P_D1_^+^Pheo^-^ to ensure efficient charge separation. At the same time, the larger near-isoenergetic antenna could also contribute to its higher luminescence, by increasing the probability of Chl_D1_^*^ decay via radiative emission with respect to photochemical re-trapping (57). This is similar to what happens in plant PSII, where the yield of photochemical trapping of excitation energy is decreased by 10-15% by the association of the Light Harvesting Complex antennas (58).

The pigment layout of Chl-f-PSII is very different from that of Chl-a-PSII and Chl-d-PSII. The 30 Chl-a molecules are energetically separated from Chl_D1_, absorbing at 720 nm, by >30 nm (>3*kB*T). This means excitation energy resides predominantly on Chl_D1_^*^ and on the other 4 far-red pigments. If the equilibration of the excitation energy between the 5 far-red pigments were significantly faster than charge separation, this pigment arrangement would result in a higher probability of populating Chl_D1_^*^ in Chl-f-PSII than in Chl-a-PSII and Chl-d-PSII (table S5). The higher Chl_D1_^*^ population in Chl-f-PSII could ensure that sufficient yield of charge separation is achieved even when the E_m_ of Phe/Phe^-^ is increased by much less that the 100 meV needed to compensate for the nominally lower energy in Chl_D1_^*^.

However, thermal equilibration of the excitation energy over the entire antenna in Chl-f-PSII might not occur due to 3 of the 4 long-wavelength antenna chlorophylls absorbing at longer wavelength than ChlD1. This type of antenna energetics could result in rapid excited state equilibration in each of the three main pigment-protein complexes (CP43, CP47 and reaction center), due to rapid energy transfer from Chl-a to Chl-f/d (with visible light excitation) followed by slower transfer from the two postulated far-red antenna pools to ChlD1, leading to a transfer-to-trap limited bottleneck. As a result, the kinetics of excitation energy transfer from the red and far-red antenna to the reaction center could be more complex than in Chl-a-PSII and Chl-d-PSII, explaining the spread in charge separation kinetics that has been suggested based on ultrafast absorption data (59) and the slower excitation energy trapping kinetics measured by time-resolved fluorescence (60).

The driving force for charge separation is decreased in Chl-f-PSII also by the smaller energy gap between Chl_D1_^*^ and P_D1_^+^Pheo^-^, estimated to be ∼ 80 meV in Chl-f-PSII compared to ∼ 160 meV in Chl-a-PSII and Chl-f-PSII (Fig. 7). This decrease in the energy gap between Chl_D1_^*^ and P_D1_^+^Pheo^-^ is necessary in Chl-f-PSII to avoid the increased photosensitivity seen in Chl-d-PSII by maintaining a large energy gap between P_D1_^+^Phe^-^ and P_D1_^+^Q_A_^-^(∼385 meV) (Fig. 7). Nonetheless, the slower excitation energy transfer and the smaller energy gap between Chl_D1_^*^ and P_D1_^+^Pheo^-^ could be partially compensated by the decreased dilution of the excitation energy on Chl_D1_^*^ arising from the small number of long-wavelength antenna pigments, resulting in only a small loss of trapping efficiency (60) and a near-negligible effect on enzyme turnover efficiency (Figures 2-4 and (5)).

This energetic balancing trick in Chl-f-PSII, which allows both reasonably high enzyme efficiency and high resilience to photodamage (by limiting recombination via the repopulation of P_D1_^+^Phe^-^) despite working with 100 meV less energy, comes with a very significant disadvantage: its absorption cross-section at long wavelength is ∼7 times smaller than that of the standard core Chl-a-antenna in visible light. In the case of Chl-f-PSII, evolution therefore seems to have prioritized the minimization of harmful charge recombination, by maintaining a big energy gap between Phe and Q_A_, over light collection and photochemical quantum efficiency. This makes sense as this system has evolved as a facultative survival mechanism, that is not advantageous when visible light is available.

In contrast, Chl-d-PSII seems to have maximized light collection at long wavelengths (with its full-size far-red antenna) and maximized the yield of charge separation (by maintaining the full Chl_D1_^*^ to P_D1_^+^Phe^-^ driving force). However, the energy shortfall at long wavelength is lost from the “energy headroom” (mainly from the transmembrane energy gap between Phe and Q_A_) that is proposed to minimize harmful charge recombination by buffering the effects of pulses of the trans-membrane electric field associated with fluctuations in light intensity (16, 61). This seems to correspond well to the shaded and stable epiphytic niche that *A. marina* occupies (7, 62–65).

Chl-d-PSII and Chl-f-PSII have evolved different strategies to do oxygenic photosynthesis in far-red light and have been impacted differently by the decrease in energy available. Understanding how the redox tuning of the electron transfer cofactors and the layout of the far-red pigments determine the trade-off between efficiency and resilience in PSII is a necessary step to inform strategies aimed at using far-red photosynthesis for agricultural and biotechnological applications.

The present findings suggest the exchange of the full Chl-a manifold to long-wavelength chlorophylls, as seen in Chl-d-PSII (*A. marina*), should allow efficient oxygenic photosynthesis, but only under constant shading and low fluctuating (stable) light conditions: e.g., for cultivation under LED light (vertical farming, etc). The more robust, facultative Chl-f PSII, would provide only a small increase in light usage efficiency due to the intrinsically low absorption cross-section in the far red, however this extension might be beneficial in a shaded canopy.

## 4 Materials and Methods

### 4.1 Cyanobacterial growth

*Acaryochloris marina* was grown in a modified liquid K-ESM medium containing 14 µM iron (66), at 30°C under constant illumination with far-red light (750 nm, Epitex; L750-01AU) at ∼30 μmol photons m^-2^ s^-1^. *Chroococcidiopsis thermalis* PCC7203 was grown in liquid BG11 medium (67) at 30°C, under two illumination conditions: white light at ∼30 μmol photons m^-2^ s^-1^ (for WL *C. thermalis* samples) and far-red light (750 nm, Epitex; L750-01AU) at ∼30 μmol photons m^-2^ s^-1^ (for FR *C. thermalis* samples). *Synechocystis sp*. PCC 6803 was grown in liquid BG11 medium at 30°C under constant illumination with white light at ∼30 μmol photons m^-2^ s^-1^. The *Thermosynechococcus elongatus ΔpsbA1, ΔpsbA2* deletion mutant (30) was grown in liquid DNT medium at 45°C.

### 4.2 Isolation of membranes and PSII cores

Membranes were prepared as described in the Supplementary Materials and Methods, frozen in liquid nitrogen and stored at −80°C until use. Partially purified *C. thermalis* PSII cores retaining oxygen evolution activity were isolated by anion exchange chromatography using a 40 ml DEAE column. The column was equilibrated with 20 mM MES-NaOH pH 6.5, 5 mM CaCl_2_, 5 mM MgCl_2_ and 0.03% (w/v) β-DM (n-Dodecyl-β-D-maltoside) and elution was done using a linear gradient of MgSO_4_ from 0 to 200 mM in 100 min (in the same buffer conditions as those used to equilibrate the column), with a flow rate of 4 ml min^-1^. Fractions enriched in PSII were pooled, frozen in liquid nitrogen and stored at −80°C. PSII-PsbA3 cores from *T. elongatus* WT*3 were purified as previously described (47).

### 4.3 Fluorescence

Flash-induced chlorophyll fluorescence and its subsequent decay were measured with a fast double modulation fluorimeter (FL 3000, PSI, Czech Republic). Excitation was provided by a saturating 70 μs flash at 630 nm and the decay in Q_A_^-^concentration was probed in the 100 μs to 100 s time region using non-actinic measuring pulses following a logarithmic profile as described in (21). The first measuring point was discarded during the data analysis because it contains a light artefact arising from the tail of the saturating flash used for excitation. Details on the analysis of the fluorescence curves are provided in the Supplementary Materials and Methods. Membrane samples were adjusted to a total chlorophyll concentration of 5 μg Chl ml^-1^ in resuspension buffer, pre-illuminated with room light for 10 seconds and then kept in the dark on ice until used for measurements. Measurements were performed at 20°C. Where indicated, 20 µM DCMU (3-(3,4-dichlorophenyl)-1,1-dimethylurea) was used.

### 4.4 Thermoluminescence and luminescence

Thermoluminescence curves and luminescence decay kinetics were measured with a laboratory-built apparatus, described in (68). Membrane samples were diluted in resuspension buffer to a final concentration of 5 µg Chl ml^-1^ in the case of *A. marina* and FR *C. thermalis* and of 10 µg ml^-1^ in the case of WL *C. thermalis* and *Synechocystis*. The samples were pre-illuminated with room light for 10 seconds and then kept in the dark on ice for at least one hour before the measurements. When used, 20 µM DCMU was added to the samples before the pre-illumination step. Excitation was provided by single turnover saturating laser flashes (Continuum Minilite II, frequency doubled to 532 nm, 5 ns FWHM). Details on the measurement conditions and on the analysis of the luminescence decay kinetics are provided in the Supplementary Materials and Methods.

### 4.5 Oxygen evolution and consumption rates

Oxygen evolution and consumption rates were measured with a Clark-type electrode (Oxygraph, Hansatech) at 25°C. Membrane samples were adjusted to a total chlorophyll concentration of 5 μg Chl ml^-1^. Illumination was provided by a white xenon lamp filtered with a heat filter plus red filter, emitting 600-700 nm light at 7100 µmol photons m^-2^ s^-1^ (Quantitherm light meter, Hansatech). When required, the light intensity was reduced by using neutral density filters (Thorlabs). For PSII maximal oxygen evolution rates, 1 mM DCBQ (2,5-Dichloro-1,4-benzoquinone) and 2 mM potassium ferricyanide were used as an electron acceptor system. Photoinhibitory illumination was performed at room temperature for 30 min with a laboratory-built red LED (660 nm, 2600 µmol photons m-^2^ s^-1^). For histidine-mediated chemical trapping of singlet oxygen, 20 µM DCMU, 5 mM L-Histidine and, where specified, 10 mM sodium azide (NaN_3_) were used. PSI activity was measured as the rate of oxygen consumption in presence of 20 µM DCMU and 100 µM methyl viologen using 5 mM sodium ascorbate and 50 µM TMPD (N,N,N′,N′-tetramethyl-p-phenylenediamine) as electron donors.

### 4.6 Flash-dependent oxygen evolution with Joliot electrode

Time-resolved oxygen polarography was performed using a custom-made centrifugable static ring-disk electrode assembly of a bare platinum working electrode and silver-ring counter electrode, as previously described (69). For each measurement, membranes equivalent to 10 µg of total chlorophyll were used. Three different light sources were used to induce the S-state transitions: a red LED (613 nm), a far-red LED (730 nm) and a Xenon flashlamp. Details on the experimental setup and on the lights used are provided in the Supplementary Materials and Methods. Measurements were performed at 20°C. For each measurement, a train of 40 flashes fired at 900 ms time interval was given and the flash-induced oxygen-evolution patterns were taken from the maximal O_2_ signal of each flash and fitted with an extended Kok model as described in (28).

### 4.7 UV transient absorption

UV pump-probe absorption measurements were performed using a lab-built Optical Parametric Oscillator (OPO)-based spectrophotometer (70) with a time resolution of 10 ns and a spectral resolution of 2 nm (see Supplementary Materials and Methods for details on the setup). Samples were diluted in resuspension buffer to a final concentration of 25 µg Chl ml^-1^ for isolated *C. thermalis* and *T. elongatus* PSII cores and 40 µg Chl ml^-1^ for *A. marina* membranes. Samples were pre-illuminated with room light for 10 seconds and then kept in the dark on ice for at least one hour before the measurements. 100 µM PPBQ (Phenyl-p-benzoquinone) was added just before the measurements. The sample was refreshed between each train of flashes. For each measurement, a train of 20 flashes (6 ns FWHM) fired at 300 ms time interval was given, and absorption changes measured at 100 ms after each flash.

## Acknowledgements

This work was supported by BBSRC grants BB/R001383/1, BB/V002015/1 and BB/R00921X. Julien Sellés acknowledges funding from the Labex Dynamo (ANR-11-LABX-0011-01). AB was in part supported by the French Infrastructure for Integrated Structural Biology (FRISBI) ANR-10-INBS-05. SS acknowledges support from Fondazione Cariplo, project “Cyanobacterial Platform Optimised for Bioproductions” (ref. 2016-0667).

## Authors contributions

A.W.R., S.V. and A.F. conceived the study; S.V. performed the fluorescence, (thermo)luminescence, luminescence and oxygen evolution measurements and analyzed the data together with A.F. and A.W.R.; W.R. performed the photoinhibition measurements and analyzed the data; D.N. and R.A. performed the polarography measurements and analyzed the data with the help of H.D.; J.S., A.B. and S.V. performed the UV transient absorption measurements and analyzed the data; S.S. contributed to data analysis and interpretation; S.V. and A.W.R. wrote the manuscript, with contributions from A.B., R.A., S.S., H.D. and A.F.. All authors approved the manuscript.

