## Supplementary figures and text for "Impact of energy limitations on function and resilience in long-wavelength Photosystem II"

**This PDF files includes:**

Supplementary text
Supplementary data: Figures S1-S14
Tables S1-S5

References for SI citations

### Supplementary materials and methods

*Isolation of membranes*

Cells were harvested by centrifugation at 6,000 x g for 5 min and resuspended in ice-cold buffer (50 mM MES-NaOH pH 6.5, 5 mM $CaCl_2$, 10 mM $MgCl_2$, 1.2 M betaine and 20% v/v glycerol) with a protease inhibitor mixture (1 mM aminocaproic acid, 1 mM benzamidine, and 0.2 % (w/v) bovine serum albumin) and 0.5 mg $ml^{-1}$ DNaseI. All following steps were performed on ice under dim green light. *A. marina* and *C. thermalis* cells were broken by two passages through a cell disruptor (Constant System, Model T5) at a pressure of 25 kPsi. *Synechocystis* cells were broken with bursts of vortexing with glass beads. Unbroken cells were removed by centrifugation for 5 min at 1,000 x g, 4ºC. Membranes were pelleted by centrifugation at 125,000 x g and 4ºC for 30 min and washed three times with resuspension buffer. Membranes were resuspended in resuspension buffer, frozen in liquid nitrogen and stored at -80°C.

*Removal of Mn-cluster by Tris-washing of membranes*

*A. marina* membranes were diluted in ice-cold 1 M Tris pH 9.5 plus 3 mM EDTA to a final chlorophyll concentration of 190 µg $ml^{-1}$ and incubated on ice under ambient light with continuous stirring for 30 min at 4°C. The membranes were then washed twice in ice-cold resuspension buffer (the same used for membrane isolation) and finally resuspended in the same.

*Analysis of $Q_A^-$ reoxidation kinetics as measured by fluorescence*

The flash-induced chlorophyll fluorescence curves were fitted with a linear combination of two exponentials (fast and middle phase) and a hyperbolic component (slow phase), where Ft is the variable fluorescence yield, $F_0$ is the basic fluorescence level before the flash, $A_1$–$A_3$ are the amplitudes and $T_1$–$T_3$ are the time-constants, based on (1, 2).

$$F_t - F_0 = A_1 \cdot \exp(-t/T_1) + A_2 \cdot \exp(-t/T_2) + A_3/(1 + t/T_3) \qquad \text{Eq.1}$$

In order to better resolve the µs to ms components associated with forward electron transfer from $Q_A^-$ to $Q_B$ or $Q_B^-$, the same curves but truncated at 1 s were fitted using a three exponentials decay and am off-set ($y_0$) accounting for the non-decaying signal in the time-window:

$$F_t - F_0 = A_1 \cdot \exp(-t/T_1) + A_2 \cdot \exp(-t/T_2) + A_3 \cdot \exp(-t/T_3) + y_0 \qquad \text{Eq.2}$$

The curves obtained in presence of 20 µM DCMU (3-(3,4-dichlorophenyl)-1,1-dimethylurea) could be fitted with two phases (one exponential and one hyperbolic) for *A. marina* and three phases (two exponentials and one hyperbolic) for WL and FR *C. thermalis*, because of the presence in both types of *C. thermalis* samples of a small initial fast phase, which probably corresponds to a small fraction of PSII centers where DCMU did not bind, as previously suggested (3).

*Thermoluminescence and luminescence*

For the $S_2Q_A^-$ and $S_2Q_B^-$ TL measurements, samples were cooled to -20°C and excited with a single turnover saturating laser flash (Continuum Minilite II, frequency doubled to 532 nm, 5 ns FWHM). The samples were then incubated in the dark at -20°C for 30 s, before heating from -20°C to 80°C at 1°C $s^{-1}$. The amplitudes of the TL peaks were normalized on the basis of the maximal oxygen evolution rates measured for each sample. For the measurement of the flash-dependence of TL, the samples were cooled to 4°C and excited with a single or multiple saturating laser flashes fired at 1 s time intervals. Samples were then heated from 4°C to 80°C at 1°C $s^{-1}$.

$S_2Q_A^-$ luminescence decay measurements were performed at a constant ($\Delta T < 0.2$°C) temperature of either 10, 20 or 30°C in presence of 20 µM DCMU. The samples were pre-equilibrated for 10 s in darkness at the given temperature before being excited with a single turnover saturating laser flash. Luminescence was then recorded from 570 ms to 300 s after the flash. The total luminescence emission was calculated as the integrated area below the decay curves normalized on the basis of the maximal oxygen evolution rates measured for each sample. The measured curves were fitted with a linear combination of three exponential components where L is the luminescence, $A_1$–$A_3$ are the amplitudes and $T_1$–$T_3$ are the lifetimes.

$$L(t) = A_1 \cdot \exp(-t/T_1) + A_2 \cdot \exp(-t/T_2) + A_3 \cdot \exp(-t/T_3) \qquad \text{Eq.3}$$

The average decay lifetime was calculated from the exponential components 2 and 3 as follows:

$$\tau_{av} = \sum_i A_i T_i / \sum_i A_i \qquad \text{Eq.4}$$

The contribution of each luminescence decay component to the total luminescence emission was calculated as

$$L_i = A_i T_i / \sum_i A_i \cdot T_i \qquad \text{Eq.5}$$

*UV transient absorption*

In the UV pump-probe absorption measurements performed using a lab-built Optical Parametric Oscillator (OPO)-based spectrophotometer, the single-turnover excitation flashes were provided by a Nd:YAG laser (Surelite II, Amplitude Technologies) at 532 nm, which pumped an OPO (Surelite OPO plus) producing monochromatic saturating flashes (6 ns FWHM) at the indicated wavelengths. The power of the flashes at the wavelengths used, measured at the level of the laser output, was: 2.7 mJ at 680 nm, 2.7 mJ at 720 nm, 3.8 mJ at 727 nm, 3.3 mJ at 734 nm, 3.7 mJ at 737 nm, 4 mJ at 749 mJ. The optics components between the laser output and the cuvettes containing the sample induce the same attenuation at all wavelengths. When indicated, the flash intensity was attenuated by 17%

using a metal grid. Detecting flashes were provided by an OPO (Horizon OPO, Amplitude Technologies) pumped by a frequency tripled Nd:YAG laser (Surelite II, Amplitude Technologies), producing monochromatic flashes (291 nm, 2 nm full-width at half-maximum) with a duration of 5 ns. The time delay between the laser delivering the excitation flashes and the laser delivering the detecting flashes was controlled by a digital delay/pulse generator (DG645, Stanford Research). The light-detecting photodiodes were protected from transmitted and scattered actinic light and fluorescence by BG39 Schott (Mainz, Germany) filters.

*Flash-dependent oxygen evolution with Joliot electrode*

For each measurement, membranes equivalent to 10 µg of total chlorophyll, brought to 750 µl with buffer A (150 mM NaCl, 25 mM MES, 1 M glycine betaine, 5 mM $MgCl_2$, and 5 mM $CaCl_2$, pH 6.2) were deposited on the electrode assembly, which was then centrifuged in a swing-out rotor at 10,000 × g for 10 min (at 4 °C). Using a home-built potentiostat, which provided an electrode polarization of -0.95 V (switched on 15 s before the first excitation flash), the current signal was recorded 20 ms before and 480 ms after each light flash, for a total of 40 flashes with a flash-spacing of 900 ms. The current signal reflects the $O_2$ reduction process at the bare platinum electrode. Three different light sources were used to induce the S-state transitions: a custom-made LED flashing device with two changeable high-performance LEDs (Luminus) and a Xenon flashlamp (EG&G Optoelectronics). The LEDs had emission peaks in the red and far-red (613 nm and 730 nm respectively) and the flashlamp was equipped with 570 nm cut-off filter suppressing shorter wavelengths and thereby photoelectric artefacts. The total energy per light flash was determined with a 1 $cm^2$ power meter (Ophir Photonics) at the exit of the light guide, which conveyed the light to the sample. The energy of the LED flashes (40 µs FWHM) was 270 µJ for the red LED and 210 µJ for the far-red LED, whereas for the flashlamp pulse (10 µs FWHM) it was 540 µJ. During the data acquisition the sample was kept at 20 °C using a Peltier and monitored by a temperature sensor immersed in the sample buffer.

### Supplementary material on fluorescence decay kinetics (section 2.1 of main text)

| **No addition (1 s)**[a] | | | |
|---|---|---|---|
| **Strain** | **Fast phase** $T_1$/Amp (ms/%) | **Middle phase** $T_2$/Amp (ms/%) | **Slow phase** $T_3$/Amp (s/%) |
| *A. marina* | 0.58±0.21 / 26±5 | 4.9±1.3 / 32±5 | — / 42±3** |
| WL *C. thermalis* | 0.50±0.09 / 32±3 | 3.7±0.4 /37 ±4 | — / 31±2 |
| FR *C. thermalis* | 0.53±0.16 / 26±4 | 4.7±0.7 / 45±4 | — / 30±3 |
| **DCMU (100s)**[b] | | | |
| **Strain** | **Not bound** $T_1$/Amp (ms/%) | **Middle phase** $T_2$/Amp (s/%) | **Slow phase** $T_3$/Amp (s/%) |
| *A. marina* | — / — | 0.98±0.58 / 19±8 | 6.5±1.0 / 81±8 |
| WL *C. thermalis* | 2.0±0.9 / 5±1 | 0.25±0.04 /17±1 | 6.9±0.3 / 78±1 |
| FR *C. thermalis* | 2.7±0.9 /6±1 | 1.31±0.35** / 14±3 | 10.4±0.8** / 80±3 |

Table S1. Time constants and relative amplitudes (%) of the different phases of fluorescence decay obtained by fitting the data in Fig. 2. Statistically significant differences according to Student's t-tests are indicated with asterisks **$p \leq 0.01$).

[a] The decay kinetics measured over 100 s in samples with no additions were truncated at 1 s and fitted with a three exponential equation allowing $y_0$ to account for the part decaying in >1 s.

[b] The data recorded in the presence of DCMU over a period of 100 s were fitted with two exponentials (only one in the case of *A. marina*) and one hyperbole.

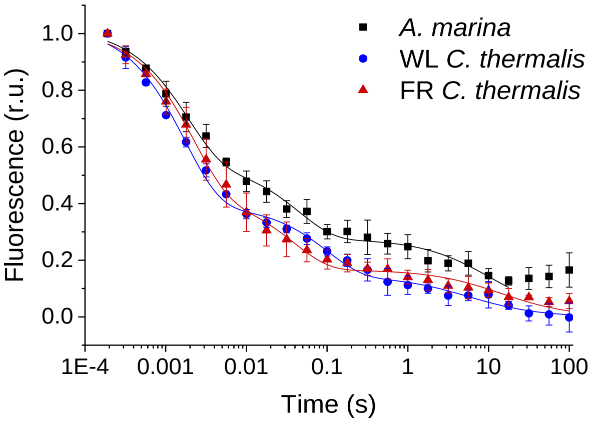


Fig. S1. Fluorescence decay kinetics after a short saturating light pulse in isolated membranes of *A. marina*, WL *C. thermalis* and FR *C. thermalis*. These are the same traces as in Fig. 2A but here including all points up to 100 s. The datapoints represent the averages of three biological replicates, ± s.d., while the lines represent the fits of the experimental data. All traces are normalized on the initial variable fluorescence ($F_m$-$F_0$, with $F_m$ measured 190 µs after the saturating flash).

| No addition 100 s | | | |
|---|---|---|---|
| | **Fast phase** | **Middle phase** | **Slow phase** |
| **Strain** | $T_1$/Amp (ms/%) | $T_2$/Amp (ms/%) | $T_3$/Amp (s/%) |
| *A. marina* | 1.8±0.3 / 47±3*** | 44.7±11.2 / 26±3 | 10.8±2.6* / 27±1**** |
| WL *C. thermalis* | 1.7±0.2 / 62±2 | 99.8±23.5*/ 24±2 | 5.6±2.4 / 14±2 |
| FR *C. thermalis* | 2.2±0.3 / 58±3 | 38.7±10.3 / 26±3 | 14.3±4.6* / 16±1 |

Table S2. Time-constants and relative amplitudes of the different phases of fluorescence decay obtained by fitting the data in Figure S1. The decay kinetics recorded over a period of 100 s were fitted with two exponentials and one hyperbole. In the case of *A. marina*, fitting of the fluorescence decay kinetics in Fig. S1 were done by excluding the datapoints between 30 and 100 s after flash, because of the presence of a non-decaying fluorescence that likely arises from a fraction of centers devoid of an intact Mn-cluster. Statistically significant differences according to Student's t-tests are indicated with asterisks (*p ≤ 0.05, ***p ≤ 0.001, ****p≤ 0.0001).

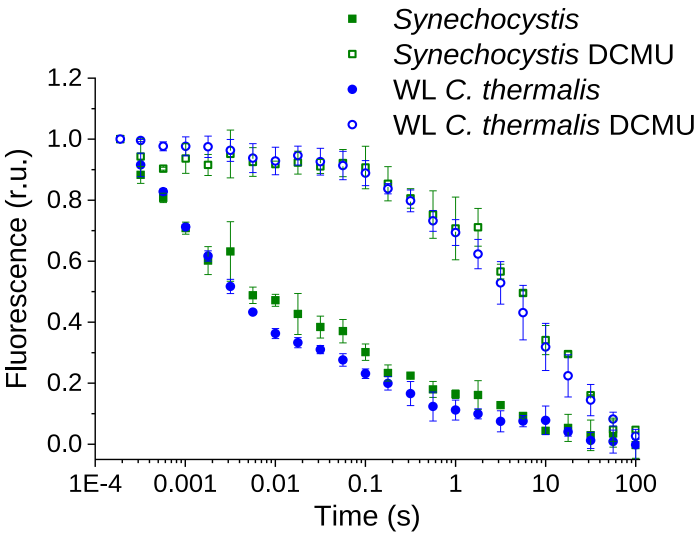


Fig. S2. Fluorescence decay kinetics after a short saturating light pulse in isolated membranes of *Synechocystis* and WL *C. thermalis* in absence and presence of DCMU. The WL *C. thermalis* data are the same as those in Fig. 2 and S1. The *Synechocystis* datapoints represent the averages of two biological replicates, ± s.d.. All traces are normalized on the initial variable fluorescence ($F_m$-$F_0$, with $F_m$ measured 190 µs after the saturating flash).

The fluorescence decay kinetics measured here in *Synechocystis* membranes, as well as those measured in *A. marina* and *C. thermalis* membranes, are faster than those measured in *Synechocystis* intact cells in previous works (4). Additionally, a study of fluorescence decay times was previously reported comparing $Q_A^-$ lifetimes in *A. marina* and *Synechocystis* but in cells rather than membranes. In *A. marina* cells the forward ($Q_A^-$ to $Q_B$) electron transfer rate was slower than in *Synechocystis* cells, while the $S_2Q_A^-$ recombination rate *A. marina* cells was faster than in *Synechocystis* cells (5). In both organisms, the fluorescence decay kinetics were faster than the values measured here in membranes. The faster rates in cells compared to isolated membranes are intrinsic to the type of sample used. The transmembrane electric field, which is present in cells but not in isolated membranes, is known to accelerate $Q_A^-$ decay both in the absence (6) and presence of DCMU (7). Additionally, the faster rates for $Q_A^-$ to $Q_B$ electron transfer in cells may be attributed to the $Q_B$ site in living cells functioning optimally at higher pH rather than at the pH 6.5 used here to maintain PSII donor-side function.

**Supplementary material on the S-state turnover efficiency (section 2.2 of main text)**

*Flash dependence of thermoluminescence.*

Fig. S3 shows the TL emission after a series of saturating flashes in *A. marina* (panels A, D and G), WL *C. thermalis* (panels B, E and H) and FR *C. thermalis* (panels C, F and I) membranes. Although no major differences in the flash patterns could be observed between the three samples, the flash dependence of the TL peak intensities (panels A, B and C) and their peak temperatures (panels D, E and F) showed variability between biological replicates. Representative TL glow curves obtained in one biological replicate for each sample after 1 to 6 flashes are shown in Fig. S3G-I. The differences in the flash patterns between replicates are easily explained by a variability in both the $S_0/S_1$ and $Q_B/Q_B^-$ ratios present in the dark before the first flash (8).

For WL *C. thermalis,* the smaller TL amplitude makes the peak temperature more difficult to estimate very precisely. For FR *C. thermalis*, a progressive broadening of the TL peak with increasing flash number made quantification less reliable, and for *A. marina* an increase in the baseline at high temperatures (also occurring to a smaller extent but still visible in WL *C. thermalis*) added to the difficulties in estimating the area of the TL peaks. For these reasons, the TL data are not precise enough to quantify potential differences in the S-state turnover efficiency in the different types of PSII, although they show that any such differences, if present, must be small (from the data in Fig. S3A, B and C).

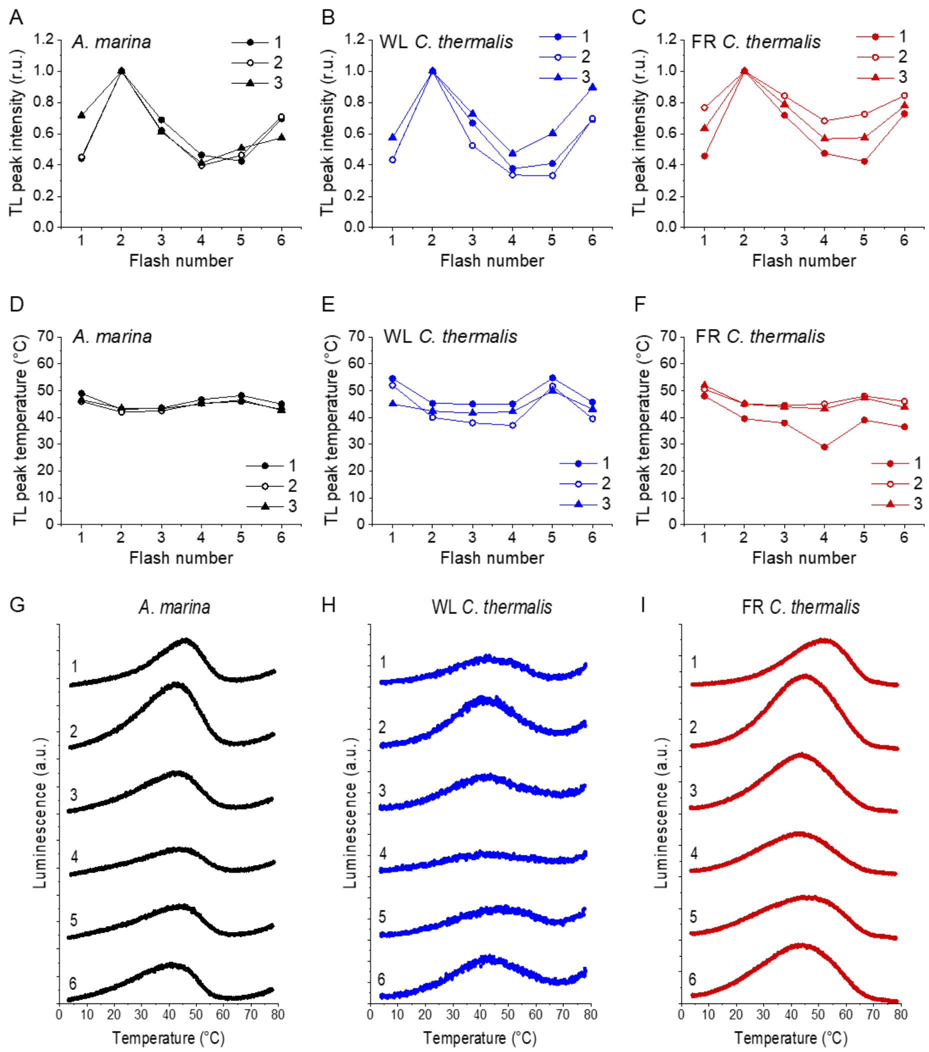


Fig. S3. Plots of the flash-induced oscillations of the thermoluminescence peak amplitudes (A, B and C) and temperatures (D, E and F) measured in *A. marina*, WL *C. thermalis* and FR *C. thermalis* membranes. The TL peak amplitudes and temperatures are plotted as a function of the number of flashes given before measuring the thermoluminescence glow curve. Peak amplitudes were normalized to the amplitude value measured after 2 flashes. The membranes, at a final concentration of 5 µg Chl ml$^{-1}$, were pre-illuminated for ~10 s at room temperature and subsequently dark-adapted on ice for 1 h before the measurements. The flashes were fired at 4°C at 1 s time intervals, and the samples were then heated from 4 to 80°C at 1°C s$^{-1}$. Each series of data points corresponds to the TL amplitudes and temperatures measured in an independent biological replicate (numbered 1 to 3). (G, H and I) Representative thermoluminescence glow curves recorded after a train of flashes (from 1 to 6, as indicated by the number next to each curve) in one of the three membrane samples in panels A-F for each strain.

*Flash-induced S-state turnover measured in the UV.*

Figure S4 shows the fit of the flash-induced absorption changes at 291 nm that reflect the progression through the S-states of the Mn-cluster (9, 10). The data are those reported in Fig. 4: absorption changes measured in *T. elongatus* PsbA3-PSII cores with excitation at 680 nm, in *A. marina* membranes with excitation at 680 nm and in partially purified Chl-f-PSII cores from FR *C. thermalis* with excitation at 680 and 750 nm. The measurements were performed in presence of PPBQ, with intervals of 300 ms between the flashes.

The fit was done by taking the absorption changes corresponding to the $S_0 \rightarrow S_1$, $S_1 \rightarrow S_2$ and $S_2 \rightarrow S_3$ transitions determined in *T. elongatus* with the procedure established by Lavergne (11), and multiplying them by a factor $\gamma$ which corresponds to the ratio in active PSII per chlorophyll of the given sample with respect to the *T. elongatus* sample. It is of note that the factor $\gamma$ indicates the fraction of active PSII centers over the total PSII present only when comparing isolated cores, while in the data reported here, in which partially purified $O_2$-evolving Chl-f-PSII cores and *A. marina* membranes were used, it merely reflects the amounts of active PSII present in those samples for a given chlorophyll concentration. Since the measurements were done by using single-turnover excitation flashes (6 ns FWHM), the double-hit parameter $\beta$ was considered to be zero. Using the formula developed by Lavorel (10), the miss parameter $\alpha$ and the proportion of the centers in $S_1$ state in the dark-adapted samples could be calculated. In these fits, the absorption changes on the first flash of the sequence was not taken into account because they may contain a non-oscillating component (11). The misses were comparable in all samples (~10%), and in the Chl-f-PSII cores from FR *C. thermalis* they did not significantly increase when using 750 nm excitation flashes.

The fits in Fig. S4 indicate that the proportion of centers in $S_1$ in the dark-adapted samples was 75% in the *A. marina* and FR *C. thermalis* samples but 100% in the *T. elongatus* PsbA3-PSII cores. All samples were pre-illuminated in ambient light for ~10s and then dark-adapted for >1 hour before the measurements and were therefore expected to be in 100% S1 at the start of the flash sequence, with ~75% of the centers having $TyrD^{\bullet}$ and ~25% having TyrD (12). It has been shown that TyrD can reduce the $S_2$ and $S_3$ oxidation states of the Mn-cluster (13): in samples having starting populations of 75% $TyrD^{\bullet}S_1$ and 25% $TyrDS_1$, some of the $S_2$ and $S_3$ states generated during the flash sequence will be re-reduced to $S_1$ and $S_2$, respectively, in centers where TyrD is present. This process will result in the apparent presence of 25% $S_0$ in the dark-adapted sample. This effect has been shown to depend on the spacing between excitation flashes (14): if the time between the flashes is not long enough to allow for TyrD donation, the flash pattern will reflect the initial presence of 100% $S_1$ (12). At room temperature, electron donation from TyrD is slower in *T. elongatus* PSII than in plant PSII (15): this could reflect the fact that *T. elongatus* is a thermophile and mesophilic cyanobacterial species such as *A. marina* and *C. thermalis* could be expected to have TyrD oxidation kinetics more similar to plants, thus explaining the difference in $S_1$ populations in our fits.

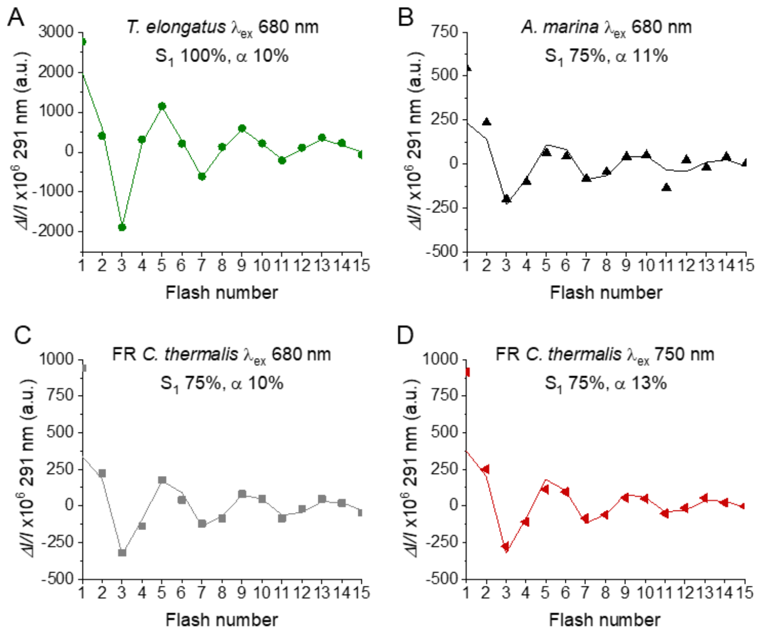


Fig. S4. Fits of the flash-induced S-state turnover measured as absorption changes at 291 nm in *T. elongatus* PsbA3-PSII cores (A), *A. marina* membranes (B) and FR *C. thermalis* PSII cores (C) with laser excitation at 680 nm and in FR *C. thermalis* PSII cores with laser excitation at 750 nm (D). Absorption changes were measured at 100 ms after each of a series of saturating flashes fired with a 300 ms time interval. The data are the same as those reported in Fig. 4, while the lines represent the fits of the experimental data. The initial fraction of PSII in $S_1$ state and the miss factors are indicated (in %).

**Supplementary material on thermoluminescence (section 2.3 of main text)**

Fig. S5 shows the plots of the peak amplitudes and temperature of the thermoluminescence arising from $S_2Q_B^-$ and $S_2Q_A^-$ in the three types of PSII. As mentioned in the main text, although our data fit qualitatively with earlier reports (5, 16), there is a degree of variability in both amplitude and temperature between biological replicates. Consequently, the average values reported in Table S3 present relatively high standard deviations. The variability in TL intensity between different membrane samples could depend on differences in the $Q_B/Q_B^-$ ratios and distribution of S states present in the dark before applying the single-turnover flash (8).

These variabilities between biological replicates could also partially explain slight discrepancies between the data reported here and those in (16) regarding the ratio of luminescence intensity between the Chl-f-PSII and the Chl-a-PSII. In Nürnberg et al. (16) the luminescence from both $S_2Q_B^-$ and $S_2Q_A^-$ were reported to be >25 times higher in FR *C. thermalis* membranes than in WL *C. thermalis* membranes, while the data reported here indicate that the luminescence of FR *C. thermalis* is between 5 and 16 times higher than in WL *C. thermalis* in the case of the $S_2Q_B^-$ recombination, and between 3 and 15 times higher in the case of the $S_2Q_A^-$ recombination. In the present work all measurements were performed at constant chlorophyll concentrations (5 µg Chl ml$^{-1}$ in the case of *A. marina* and FR *C. thermalis*, 10 µg ml$^{-1}$ in the case of WL *C. thermalis*), while in Nürnberg et al. the FR *C. thermalis* membranes were diluted to achieve a signal intensity comparable to that obtained in WL *C. thermalis* membranes. Although the dilution factor was included in the normalization on the $O_2$ evolution activities, these differences in the protocols used could contribute to the quantitative discrepancies, together with the biological variability, as at higher chlorophyll concentrations sample self-absorption can occur, thus skewing the measured TL intensity.

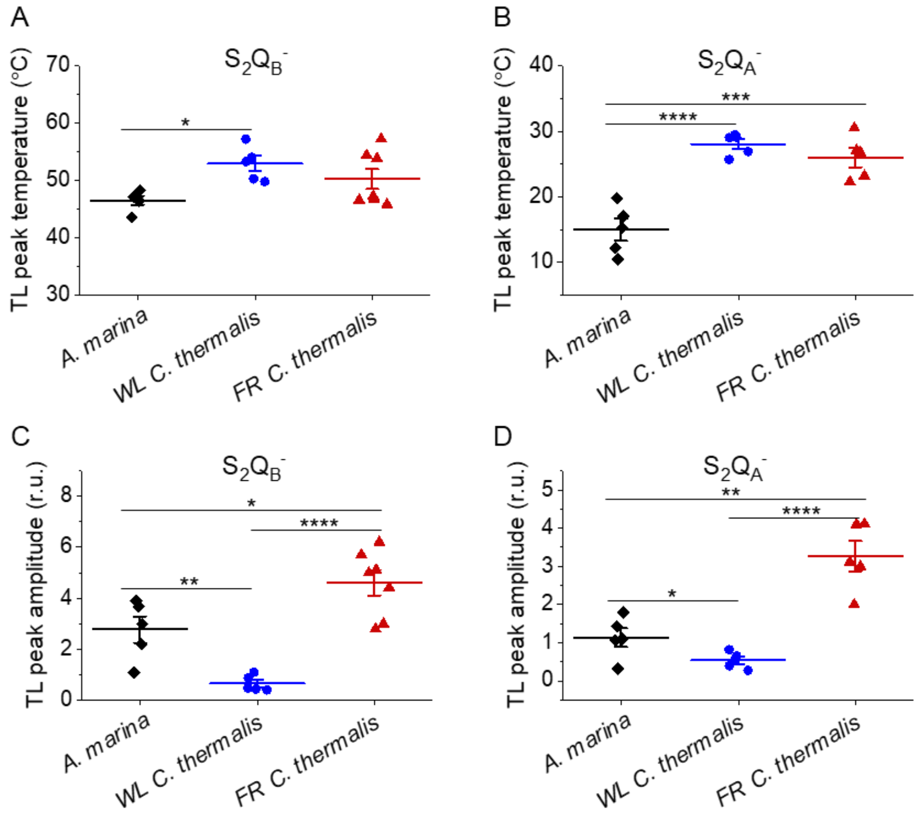


Fig. S5. Thermoluminescence in *A. marina*, WL *C. thermalis* and FR *C. thermalis* membranes. Plots of the temperatures (A and B) and of the normalized amplitudes (C and D) of the thermoluminescence peaks deriving from $S_2Q_B^-$ and from $S_2Q_A^-$ back-reaction, including the examples shown in Fig. 5A and B. Each point represents an independent biological replicate, the horizontal lines represent the mean values, ± standard error (for $S_2Q_B^-$: *A. marina* n=5, WL *C. thermalis* n=5, FR *C. thermalis* n=7; for $S_2Q_A^-$: *A. marina* n=5, WL *C. thermalis* n=5, FR *C. thermalis* n=5). Statistically significant differences according to Student's t-tests are indicated with asterisks (*$p \leq 0.05$, **$p \leq 0.01$, ***$p \leq 0.001$, ****$p \leq 0.0001$).

| Strain | $S_2Q_B^-$ | | $S_2Q_A^-$ | | ΔT (°C) |
|---|---|---|---|---|---|
| | T (°C) | Amp (r.u.) | T (°C) | Amp (r.u.) | |
| *A. marina* | 46.5±1.8 | 2.77±1.15 | 14.9±3.7 | 1.15±0.55 | 31.5±2.8* |
| WL *C. thermalis* | 52.9±3 | 0.65±0.31 | 28.1±1.7 | 0.54±0.21 | 24.9±3.2 |
| FR *C. thermalis* | 50.3±4.7 | 4.61±1.29 | 26±3.3 | 3.27±0.88 | 24.3±5.1 |

Table S3. Average values (±s.d.) of the temperatures (T) and of the normalized amplitudes (Amp, in relative units) of the thermoluminescence peaks from $S_2Q_B^-$ and from $S_2Q_A^-$ back-reactions, plotted in Fig. S5. The difference in temperature between the $S_2Q_B^-$ and the $S_2Q_A^-$ (ΔT) is also reported. The ΔT in *A. marina* is significantly bigger than the one in WL and FR *C. thermalis* according to Student's t-test, as indicated with an asterisk (*p ≤ 0.05).

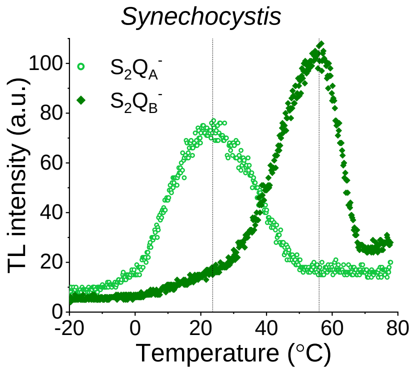


Fig. S6. Thermoluminescence measured in the absence of inhibitors ($S_2Q_B^-$) or in the presence of DCMU ($S_2Q_A^-$) in *Synechocystis* membranes. The dashed vertical lines indicate the two peak positions.

TL measurements were performed in *Synechocystis* membranes to compare the $S_2Q_A^-$ (measured in presence of DCMU) and $S_2Q_B^-$ peak temperatures with those measured in the *A. marina* and *C. thermalis* membranes under the same conditions. The TL peak temperatures in *Synechocystis* were comparable with those in WL and FR *C. thermalis*, confirming that the $S_2Q_A^-$ in Chl-d-PSII recombines at a lower temperature than in Chl-a-PSII, as previously reported in cells (3), and in Chl-f-PSII (Fig. 5 and S5).

### Supplementary material on luminescence decay (section 2.3 of main text)

The $S_2Q_A^-$ luminescence decay curves measured in *A. marina*, WL *C. thermalis* and FR *C. thermalis* at 10, 20 and 30°C (Fig. S7) could be fitted with three exponential components (Table S4) and the differences in the kinetics between samples and between temperatures could be ascribed to differences in the amplitude and lifetimes of these components.

The luminescence decays at each temperature were similar in shape in Chl-a-PSII and Chl-f-PSII, while they were markedly different in Chl-d-PSII. Chl-a-PSII and Chl-f-PSII had a fast decay phase ($T_1$ ~0.5 and ~1 s, respectively) absent in Chl-d-PSII. This phase, that has a bigger amplitude in Chl-a-PSII (~60%) than in Chl-f-PSII (~30%), is too fast to correspond to the $S_2Q_A^-$ recombination and appears to match the recombination rates for $TyrZ^{\bullet}(H^+)Q_A^-$, a reaction that dominates in centers lacking the Mn cluster (17). The contribution of this fast component to the total luminescence emission was no more than 10% in the case of WL *C. thermalis* and 5% for FR *C. thermalis*. This decay phase was not detectable in the case of *A. marina*, suggesting that $TyrZ^{\bullet}(H^+)Q_A^-$ recombination might be too fast in Chl-d-PSII to appear in our measurements. In *A. marina* an additional slower phase (~40s) was present at 10°C, but the very low amplitude made its contribution to the overall decay negligible.

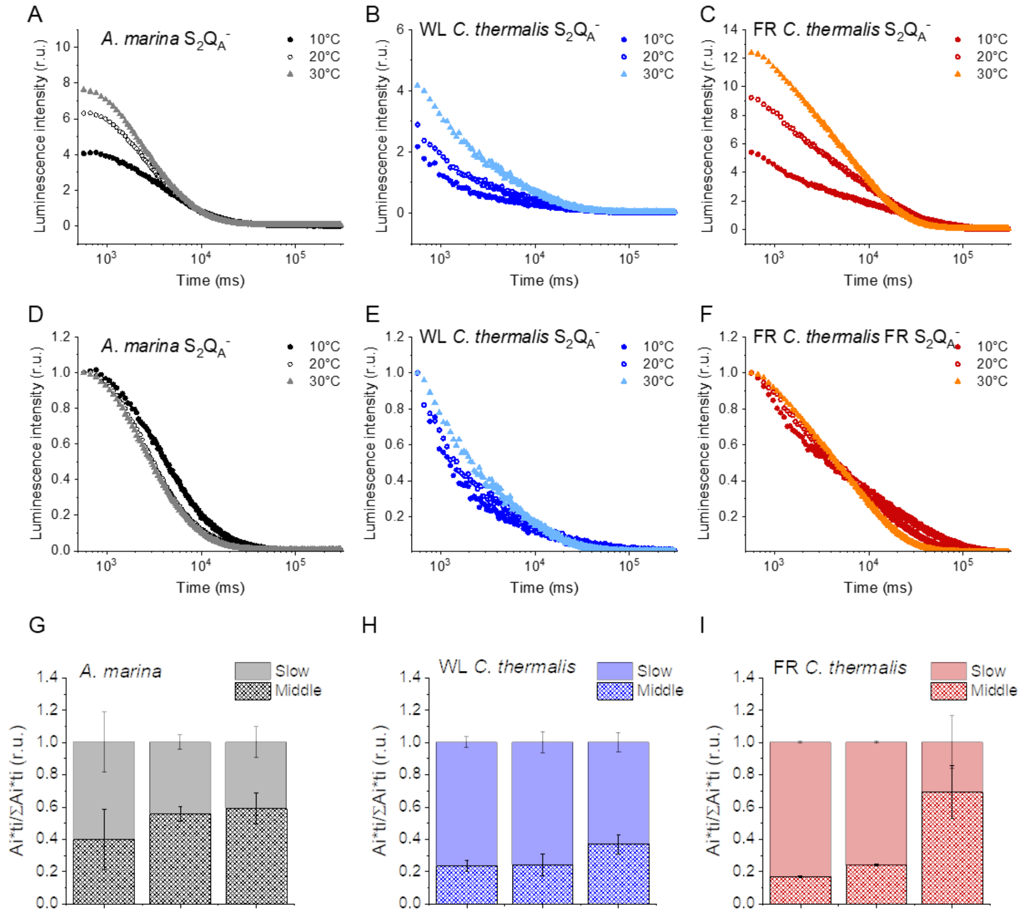


Fig. S7. (A, B and C) Representative $S_2Q_A^-$ luminescence decay curves measured in *A. marina*, WL *C. thermalis* and FR *C. thermalis* membranes in the presence of DCMU. The measurements were performed at 10, 20 and 30°C. The luminescence decays were measured for 300 s after the flash, and the time is plotted on a logarithmic scale. (D, E and F) The same curves as in (A-C) after normalization on the initial intensities. (G, H and I) Relative contributions of the middle and slow decay components to the total luminescence emission arising from $S_2Q_A^-$ recombination, calculated using the values in Table S4.

The luminescence decay that we ascribe to the $S_2Q_A^-$ back-reaction in the seconds to tens of seconds timescale, is comprised of two decay components, designated the middle and slow phases in Table S4. Both phases were faster in Chl-d-PSII (~3 and ~11 s) than in Chl-a-PSII (~4 and ~25 s), but slower in Chl-f-PSII (~9 and ~39 s). In *A. marina* the lifetimes of these two decay components did not show a significant temperature dependence, resulting in only a minor acceleration of the overall luminescence decay of the Chl-d-PSII between 10 and 30°C (Fig. 5D). Indeed, the relative contribution of the two decay phases to the total luminescence changed little in function of temperature in this sample (Fig. S7G), with the changes being only at the level of the amplitude of the decay phases. The middle phase lifetimes did not show a significant temperature dependence in WL and FR *C. thermalis* either, but they were slower than in *A. marina*, especially in FR *C. thermalis*. The slow phase was also slower in the two *C. thermalis* samples and, additionally, its decay accelerated with increasing temperature, while its amplitude decreased. This resulted in its contribution to the overall luminescence decreasing between 10 and 30°C (Fig. S7H and I) and the overall decay accelerating significantly (Fig. 5D), especially in the FR *C. thermalis*.

| **Strain and temperature** | **Fast phase** $T_1$/Amp (s/%) | **Middle phase** $T_2$/Amp (s/%) | **Slow phase** $T_3$/Amp (s/%) | **Additional phase** $T_4$/Amp (s/%) |
|---|---|---|---|---|
| ***A. marina*** | | | | |
| 10°C | — / — | 3.5±1.0 / 64±22 | 10.6±2.9 / 35±18 | 36.5±7.3 / 5±3 |
| 20°C | — / — | 3.2±0.6 / 88±5 | 12.7±2.9 / 18±3 | — / — |
| 30°C | — / — | 3.0±0.4 / 87±9 | 10.2±2.4 / 19±7 | — / — |
| **WL *C. thermalis*** | | | | |
| 10°C | 0.5±0.2 / 72±8 | 4.0±2.4 / 18±4 | 32.0±6.8 / 7±2 | — / — |
| 20°C | 0.4±0.1 / 62±13 | 3.2±1.2 / 23±7 | 19.1±2.8 / 12±5 | — / — |
| 30°C | 0.6±0.1 / 55±7 | 6.0±0.8 / 32±4 | 16.9# / 9# | — / — |
| **FR *C. thermalis*** | | | | |
| 10°C | 1.0±0.2 / 43±2 | 10.4±1.0 / 26±2 | 43.7±1.8 / 31±2 | — / — |
| 20°C | 1.0±0.1 / 28±4 | 7.9±0.7 / 35±3 | 23.5±1.8 / 38±4 | — / — |
| 30°C | 1.4±0.4 / 14±10 | 8.6±1.1 / 72±18 | 18.6±3.7 / 15±8 | — / — |

Table S4. Time constants and relative amplitudes of the different phases of luminescence decay obtained by fitting the data recorded at 10, 20 and 30°C with a three-exponential equation. The values represent the averages of 3 biological replicates, ± s.d. The fast decay phase is assigned to $TyrZ^{\bullet}(H^+)Q_A^-$ recombination, while the middle and slow phases are assigned to $S_2Q_A^-$ recombination. The additional phase identified in *A. marina* membranes at 10°C is unassigned. #The slow phase in WL *C. thermalis* membranes at 30°C could be reliably fitted only in one replicate out of three.

It can be argued that the differences in kinetics between samples and their changes in function of temperature could represent changes in the relative contribution of different recombination pathways to the decay of $S_2Q_A^-$. It is not clear though whether each of the two decay components we identified represents a distinct recombination route or whether they derive from the combination of more complex kinetics. For instance, it has been suggested that the so-called "deactivation" luminescence should follow a hyperbolic decay, rather than an exponential decay, due to the progressive decrease in the concentration of $S_2Q_A^-$ resulting in a progressive slowing down of the rates of the various recombination routes (18, 19). The data presented here could be satisfactorily fitted with exponentials but, given the considerations above and the uncertainty about how the evolution of luminescence reflects the actual concentrations of the charge separated states from which it originates, no assignment of the decay phase to specific recombination routes could be made.

Altogether, the data show that the luminescence kinetics in Chl-d-PSII are significantly different from those in Chl-a-PSII and Chl-f-PSII, pointing to a faster decay of the $S_2Q_A^-$ charge separated state.

According to electron tunnelling calculations, the rate of $P_{D1}^+Q_A^-$ direct recombination to ground ($10^2$-$10^3$ $s^{-1}$) is much slower than $P_{D1}^+Phe^-$ recombination to ground ($10^6$-$10^7$ $s^{-1}$), although the limiting rate for $S_2Q_A^-$ recombination via the repopulation of $Phe^-$ is thought to be the migration of the electron hole from the Mn-cluster to TyrZ (~$10^3$ $s^{-1}$) (20, 21). Although the temperature dependence of the recombination routes is complex, an increase in temperature would have no effect on the rate of $P_{D1}^+Q_A^-$ direct recombination to ground, but would increase the rate of the backwards electron transfer from $Q_A^-$ to Phe, as this is thermally activated, following the relationship

$$k_{rev} = k_{fwd} \cdot e^{-\frac{\Delta G^0}{k_B T}} \qquad \text{Eq. 6}$$

where $k_{rev}$ and $k_{for}$ are the rate constants of the backward and forward electron transfer, respectively, $\Delta G$ is the energy gap between the two cofactors, $k_B$ is the Boltzmann constant and T is the temperature. Note that the rates of back-transfer of the positive charge from the Mn-cluster to $P_{D1}$ are also thermally activated and thus will accelerate with temperature and affect the rates of recombination from $Phe^-$. In this case, though, the smaller $\Delta$Gs involved should result in a less pronounced temperature dependence compared to the back electron transfer from $Q_A^-$ to Phe (according to the equation above).

The acceleration of the luminescence decay kinetics with increasing temperature, observed for Chl-a-PSII and Chl-f-PSII, could reflect an increase in the contribution of $S_2Q_A^-$ recombination route via repopulation of $Phe^-$ in competition with the direct, non-radiative $P_{D1}^+Q_A^-$ recombination route. In Chl-d-PSII, the lower temperature of the $S_2Q_A^-$ recombination thermoluminescence peak (Fig. 5B and Table S3) suggests a smaller $\Delta$G between $Q_A$ and Phe. This would result in a faster electron transfer from $Q_A^-$ back to Phe, with this route already dominating the competition with the direct $P_{D1}^+Q_A^-$

recombination to ground, with a consequently high luminescence yield and a small temperature sensitivity of the decay rates.

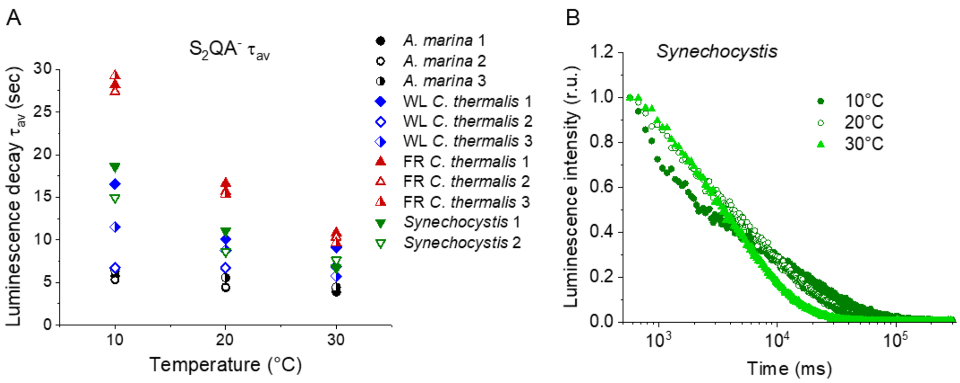


Fig. S8. (A) Plots of the average $S_2Q_A^-$ luminescence decay lifetimes ($\tau_{av}$) in *A. marina*, WL *C. thermalis*, FR *C. thermalis* and *Synechocystis* membranes. Each series of data corresponds to an independent biological replicate. The *A. marina*, WL *C. thermalis* and FR *C. thermalis* datasets are those used to calculate the average decay values plotted in Fig. 5D. (B) Representative $S_2Q_A^-$ luminescence decay curves measured in *Synechocystis* membranes at 10, 20 and 30°C, after normalization on the initial intensities. The luminescence decays were measured for 300 s after the flash and plotted on a logarithmic scale.

### Supplementary material on singlet oxygen production and sensitivity to high light in the far-red PSII (section 2.4 of main text)

*Light sources*

Given the different pigments involved in light capture in the three types of PSII studied here, the comparability of experiments could be adversely affected by differences in excitation rates due to the degree of matching of the absorption spectrum of the PSII with excitation spectrum of the light sources used. Fig. S9A shows absorption spectra of the three membrane preparations containing the 3 types of PSII and shows the spectral profiles of the xenon lamp and the 660 nm LED. For both light sources the WL and the FR *C. thermalis* samples have a greater spectral overlap with the actinic light spectrum, than does the *A. marina* sample. It can be concluded that under identical illumination conditions, *A. marina* would receive less photons during a period of illumination compared to two *C. thermalis* samples.

Fig. S9B shows the oxygen evolution in the presence of the electron acceptor system, and oxygen consumption rates in the presence of histidine measured in *A. marina* membranes as a function of the light intensity. The figure shows experiments done in three biological replicates. Both rates showed a comparable dependence on light intensity and saturated at 7100 µmol photons $m^{-2}$ $s^{-1}$, the intensity used in all the oxygen measurements. The same light intensity was saturating also in the case of WL and FR *C. thermalis* membranes used at the same concentration of 5 µg Chl ml$^{-1}$ (Fig. S9C).

Fig. S9D and E show that both the LED and the xenon lamp gave the same rates of $O_2$ evolution, and given their different actinic spectra, this indicates that both were saturating under the conditions of the experiment.

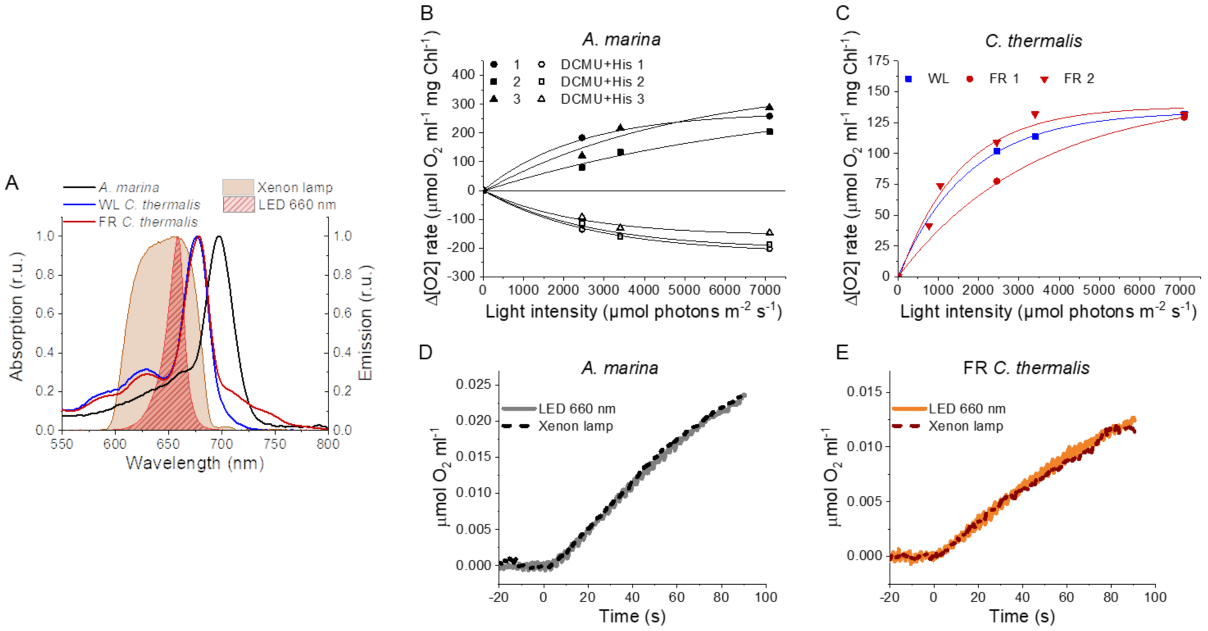


Fig. S9. Light sources used for $^1O_2$ production measurements and high light treatment. (A) Absorption spectra (normalized on the maximal absorption in the Qy region) of *A. marina*, WL *C. thermalis* and FR *C. thermalis* membranes and spectral profiles (normalized on the maximal emission) of the 660 nm LED and xenon lamp used. (B) Light saturation curves of $O_2$ evolution (in presence of DCBQ and ferricyanide, solid symbols) and $^1O_2$ production (in the presence of DCMU and histidine, open symbols) in three biological replicates of *A. marina* membranes, using the xenon lamp. The intensity of the lamp was decreased by using neutral filters. (C) Light saturation curves of $O_2$ evolution in WL *C. thermalis* (1 biological replicate) and FR *C. thermalis* (2 biological replicates) membranes, used at a final Chl concentration of 5 µg ml$^{-1}$. (D and E) Representative $O_2$ electrode traces monitoring maximal $O_2$ evolution in *A. marina* and FR *C. thermalis* membranes, used at a final Chl concentration of 5 µg ml$^{-1}$. Measurements were performed in presence of DCBQ and potassium ferricyanide using either the 660 nm LED (2600 µmol photons m-$^{2}$ s$^{-1}$) or the xenon lamp (7100 µmol photons m-$^{2}$ s$^{-1}$) for illumination.

*Singlet oxygen production in membranes*

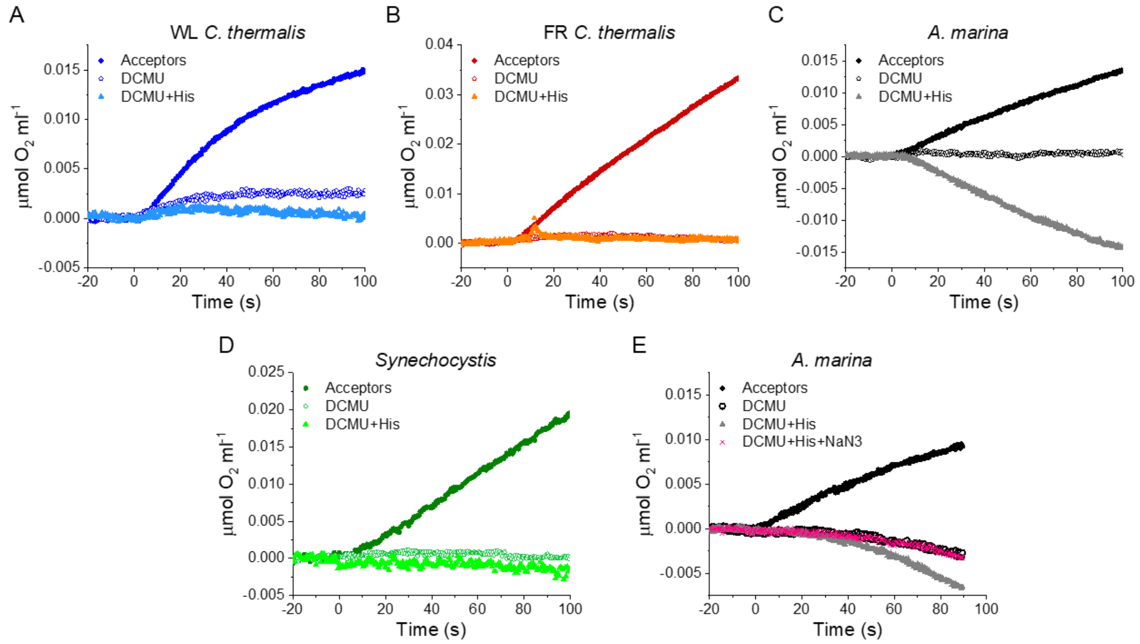


Fig. S10. Singlet oxygen production in *A. marina*, WL *C. thermalis*, FR *C. thermalis* and *Synechocystis* membranes. All samples were used at a chlorophyll concentration of 5 µg $ml^{-1}$. (A, B, C and D) Representative $O_2$ electrode traces monitoring $O_2$ evolution and uptake. $^1O_2$ production in presence of DCMU was measured as the rate of histidine-dependent consumption of $O_2$ induced by saturating illumination (xenon lamp, 7100 µmol photons $m^{-2}$ $s^{-1}$). Measurements were performed in in presence of DCBQ and ferricyanide (Acceptors), or in presence of DCMU, with or without the addition of histidine (His). (E) $^1O_2$ production in a different *A. marina* membrane preparation showing the effect of sodium azide ($NaN_3$). Sodium azide is a $^1O_2$ quencher that regenerates $O_2$ in competition with $^1O_2$ scavenging by histidine.

*Singlet oxygen production experiments: the presence of the Mn cluster.*

To test whether the $^1O_2$ production in *A. marina* was related to the fraction of PSII centers devoid of an intact Mn-cluster, which is the most obvious functional difference between *A. marina* membrane samples and those from the WL and FR *C. thermalis*, we compared $^1O_2$ formation in untreated and Tris-washed membranes. Tris-washing was used to remove the Mn-cluster from all PSII. As shown in Fig. S11, the Tris-washed membranes did not display any $O_2$ evolution activity in presence of the acceptors DCBQ and potassium ferricyanide but retained the same $^1O_2$ production capacity as the untreated sample. This indicates that $^1O_2$ formation in *A. marina* is not related to the fraction of centers that are capable of water oxidation.

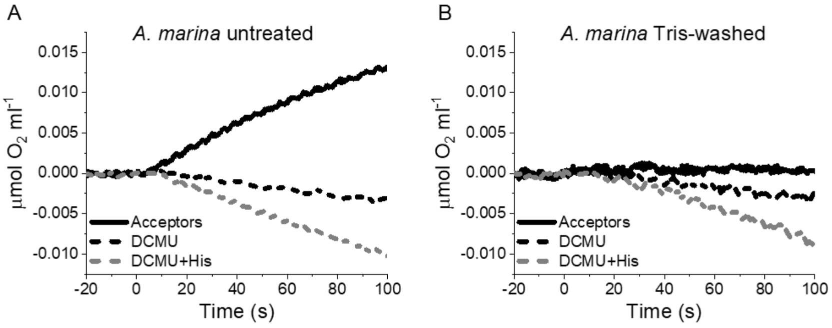


Fig. S11. [1]$O_2$ formation in presence of DCMU measured as the rate of histidine-dependent consumption of $O_2$ induced by saturating illumination in untreated (A) and Tris-washed (B) *A. marina* membranes. Measurements were performed in the presence of DCBQ and potassium ferricyanide (Acceptors) or in presence of DCMU, with or without the addition of L-Histidine (His).

*Singlet oxygen production: does PSI contribute to $O_2$ uptake?*

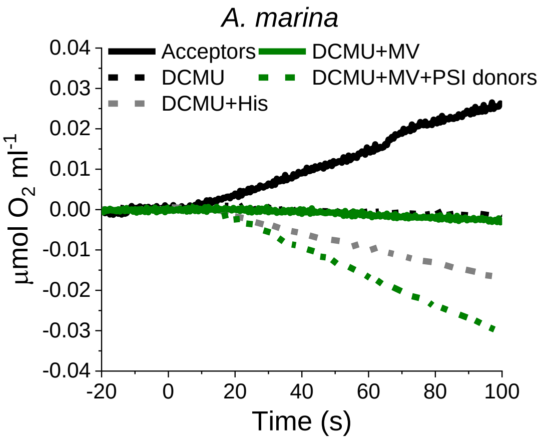


Fig. S12. $O_2$ electrode traces monitoring $O_2$ evolution and uptake in *A. marina* membranes; [1]$O_2$ formation is monitored by $O_2$-uptake due to [1]$O_2$ scavenging by histidine. Measurements were performed in the presence of DCBQ and potassium ferricyanide (Acceptors) or in the presence of DCMU, with or without the addition of L-Histidine (His). PSI activity (green traces) was measured as the rate of methyl viologen (MV, 100 μM)-dependent oxygen consumption in the presence of DCMU, either with (dashed green line) or without (solid green line, "PSI donors") the electron donors ascorbate (5 mM) and TMPD (50 μM).

We tested whether light-induced oxygen consumption observed in *A. marina* membranes could be derived from Photosystem I (PSI) turnover. It is well-known that PSI can reduce $O_2$ to $O_2^{\bullet-}$ and this is greatly enhanced by methyl viologen (MV) acting as a redox mediator. The PSI electron donors, plastocyanin or cytochrome $c_6$, which are both soluble in the lumen, are expected to be lost during preparation of the membranes. As a result, illumination is likely to accumulate oxidized $P_{700}$ resulting in PSI being non-functional. To confirm this in *A. marina* membranes in which PSII activity was blocked by DCMU, we tested whether methyl viologen (MV) could induce a light-dependent oxygen consumption in the absence of the histidine $^1O_2$ trap. In isolated *A. marina* membranes, no MV-mediated oxygen consumption was observed in presence of DCMU unless the exogenous PSI electron donors, ascorbate and TMPD, were also added (Fig. S12). This demonstrates that under the conditions of the experiments used to estimate $^1O_2$ trapping by histidine in the isolated *A. marina* membranes, there was no contribution from PSI activity.

*Singlet oxygen production in cells*

We tested if differences in the stability of the membrane samples could explain the marked increase in singlet oxygen production in *A. marina* compared to both the WL and FR the *C. thermalis* samples (see Fig. 6 and related text). Lower stability of PSII in the isolated membranes of *A. marina* was suggested by the presence of long-lived non-decaying emission observed when measuring fluorescence decay kinetics (Fig. 2 and S1) and attributed to a fraction of centers devoid of an intact Mn-cluster. We therefore used the histidine trapping method to compare the rates of singlet oxygen production in *A. marina* and FR *C. thermalis* intact cells. The reliability of the His-trapping method to monitor $^1O_2$ production in intact cyanobacterial cells has been previously demonstrated (22).

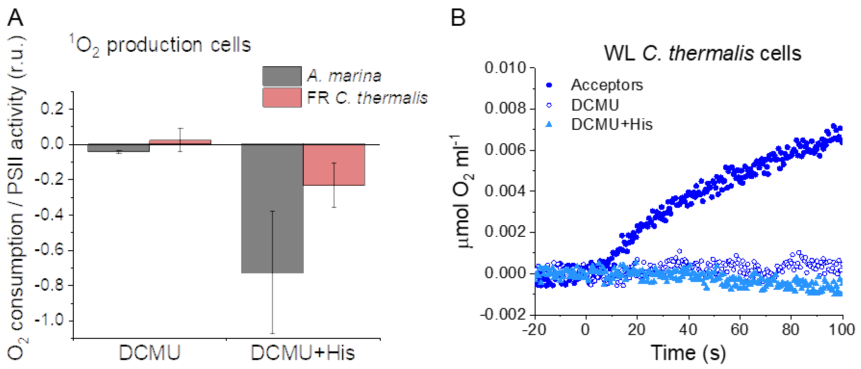


Fig. S13. Singlet oxygen production in intact cells. (A) $^1O_2$formation in presence of DCMU measured as the rate of histidine-dependent consumption of $O_2$ induced by saturating illumination in *A. marina* and FR *C. thermalis* cells. The data are averages (±s.d.) of 3 biological replicates for each strain. For each replicate, the rates of oxygen consumption were normalized to the maximal oxygen evolution rates obtained with the same illumination in the presence of the exogenous acceptors, DCBQ and ferricyanide. (B) $O_2$ electrode traces monitoring $O_2$ evolution and uptake in WL *C. thermalis* cells. $^1O_2$ production in the presence of DCMU was measured as the rate of histidine-dependent consumption of $O_2$ induced by saturating illumination (xenon lamp, 7100 µmol photons $m^{-2}$ $s^{-1}$). Measurements were performed in the presence of DCBQ and ferricyanide (Acceptors), or in the presence of DCMU, with or without the addition of histidine (His).

Fig. S13A shows that in *A. marina* cells the rate of histidine-mediated oxygen uptake was much higher, relative to the maximal oxygen evolution rate, than in FR *C. thermalis*. The values obtained in cells were comparable with those obtained in isolated membranes, despite variability between biological replicates (this variability makes the difference between the two strains less significant than that measured in membranes, p = 0.08). Like FR *C. thermalis* cells, WL *C. thermalis* cells also showed low levels of $^1O_2$ production, similar to those measured in the respective membranes (Fig. S13B). It is of note that both in membranes and intact cells, the rates of maximal $O_2$ evolution (measured in presence of exogenous electron acceptors) and of $^1O_2$ production (measured in presence of DCMU) do not depend on the functionality of the electron transport chain downstream of PSII.

**Supplementary material on the D1-Q130E occurrence in different species (section 3.2 of main text)**

```
A
T. elong PsbA1        110 GPYQLIIFHFLLGASCYMGRQWELSYRLGMRPWI 143
T. elong PsbA3        110 GPYQLIIFHFLIGVFCYMGREWELSYRLGMRPWI 143
C. therm FR           111 GPYQMIGFHYIPALCCYAGREWELSYRLGMRPWI 144
C. therm WL1          110 GPYQLVIFHFLIGCFCYMGRQWELSYRLGMRPWI 143
C. therm WL2          110 GPYQLVIFHFLIGVFCYMGREWELSYRLGMRPWI 143
C. therm WL3          110 GPYQLVIFHFLIGVFCYMGREWELSYRLGMRPWI 143
A. marin 1            113 GPYQLIILHFLIAIWTYLGRQWELSYRLGMRPWI 146
A. marin 2            110 GPYQLIIFHYMIGCICYLGRQWEYSYRLGMRPWI 143
A. marin 3            110 GPYQLIIFHYMIGCICYLGRQWEYSYRLGMRPWI 143

B
Leptol JSC-1          110 GPYQMIAAHYVPALCCYMGREWELSYRLGMRPWI 143
Oscill JSC-12         111 GPYQMIGAHYIPALACYMGRQWELSYRLGMRPWI 144
Caloth NIES-267       110 GPYQMIAFHYIPALSCYMGREWELSYRLGMRPWI 143
Mastigo BC008         111 GPYQMIAFHYIPALACYMGREWELSYRLGMRPWI 144
C. therm FR           111 GPYQMIGFHYIPALCCYAGREWELSYRLGMRPWI 144
Caloth PCC7507        111 GPYQMIAFHYIPALSCYMGREWELSYRLGMRPWI 144
Caloth NIES-3974      111 GPYQMIAFHYIPALACYMGREWELSYRLGMRPWI 144
Fische NIES-592       111 GPYQMIGFHYIPALACYMGREWELSYRLGMRPWI 144
Fische NIES-3754      111 GPYQMIGFHYIPALACYMGREWELSYRLGMRPWI 144
Mastigo SAG4.84       111 GPYQMIGFHYIPALACYMGREWELSYRLGMRPWI 144
Chlorog PCC6912       111 GPYQMIGFHYIPALACYMGREWELSYRLGMRPWI 144
Fische PCC9605        111 GPYQMIGFHYIPALACYMGREWELSYRLGMRPWI 144
Halomicr. Hongd.      110 GPYQMIAFHYIPALLCYMGREWELSYRLGMRPWI 143
Synechoco PCC7335     109 GPYQMIAFHYIPALLCYLGREWELSYRLGMRPWI 142
Pleuroc CCALA161      110 GPYQMIAFHYIPALCCYLGREWELSYRLGMRPWI 143
Hydroco NIES-593      110 GPYQMIALHYVPALCCYLGREWELSYRLGMRPWI 143
Pleuroc PCC7327       110 GPYQMIALHYVPALCCYLGREWELSYRLGMRPWI 143
```

Fig. S14. Occurrence of the high light-associated D1-Gln130Glu substitution in the different types of PSII. (A) Multi-alignment of the D1 proteins of *T. elongatus*, *C. thermalis* and *A. marina*. (B) Multi-alignment of the far-red light induced D1 isoforms of *C. thermalis* and other Chl-f species. Both alignments were done using Clustal Omega (23), the sequences were retrieved from the KEGG (https://www.kegg.jp/) and NCBI (https://www.ncbi.nlm.nih.gov/) databases. For each alignment only a 33 amino acid region is shown, the start and end positions with respect to each full sequence are indicated with numbers. The Q130E substitution is highlighted as white font on black background. The far-red D1 sequence from *C. thermalis* is framed in red.

The high light induced D1 isoform of *T. elongatus*, PsbA3, contains a glutamate in position 130, in place of the glutamine that is present in the isoform normally expressed under low-light conditions,

PsbA1. The glutamate forms a stronger H-bond with $Phe_{D1}$, thus increasing its redox potential. This substitution is present also in high-light-induced D1 isoforms of other cyanobacterial species, and is associated with photoprotection (24).

The multi-alignment in Fig. S14A shows that the same Q→E substitution is present also in the far-red light-induced D1 isoform of *C. thermalis* (*C. therm* FR) and in two out of three of its non-far-red induced D1 isoforms (*C. therm* WL2 and 3) but is not present in any of the three D1 isoforms of *A. marina*. To date, we have not yet investigated which D1 isoform is expressed in *C. thermalis* in our white light growth conditions.

The presence of E130 is conserved in the far-red light induced D1 isoforms of most of the cyanobacteria species capable of far-red light photo-acclimation (Fig. S14B).

### Supplementary material for section 3.3 of main text

| **Chl-a-PSII** | | | |
|---|---|---|---|
| State | E* | n | Pi |
| Bulk Chl-a/Pheo-a | 685 | 34 | 0.878 |
| $Chl_{D1680}$ | 685 | 1 | **0.026** |
| $F_{685}$ | 685 | 1 | 0.026 |
| $F_{695}$ | 695 | 1 | 0.071 |
| **Chl-d-PSII** | | | |
| State | E* | n | Pi |
| Chl-a/Pheo-a | 685 | 3 | 0.002 |
| $Chl_{D1720}$ | 725 | 1 | **0.029** |
| Bulk Chl-d | 725 | 33 | 0.969 |
| **Chl-f-PSII** | | | |
| State | E* | n | Pi |
| Bulk Chl-a/Pheo-a | 685 | 32 | 0.046 |
| $Chl_{D1721}$ | 726 | 1 | **0.075** |
| $F_{720}/A_{715}$ | 720 | 1 | 0.043 |
| $F_{731}/A_{726}$ | 731 | 1 | 0.117 |
| $F_{737}/A_{732}$ | 737 | 1 | 0.2 |
| $F_{748}/A_{743}$ | 748 | 1 | 0.52 |

Table S5. Excitation energy partitions calculated for the three types of PSII assuming excitation equilibration between the pigments. $E^*$ denotes the energy of the excited state, obtained by applying a +5 nm Stoke's shift to the absorption of the pigments, n is the number of pigments belonging to each state and Pi is the normalized partition of the excited states, calculated following Boltzmann distribution (25).

The states are denoted as follows: $Chl_{D1}$ is the primary donor (Pi highlighted in bold), Bulk indicates the antenna pigments considered as isoenergetic, and F indicates the antenna pigments considered separately from the bulk, with the fluorescence emission wavelength indicated. In the case of the far-red pigments in Chl-f-PSII the peak absorptions (A) are also indicated, as taken from (26).
